# Consequences of the divergence of Methionine AdenosylTransferase

**DOI:** 10.1101/2020.08.17.254151

**Authors:** Bhanupratap Singh Chouhan, Madhuri H. Gade, Desirae Martinez, Saacnicteh Toledo-Patino, Paola Laurino

## Abstract

Methionine adenosyltransferase (MAT), which catalyzes the biosynthesis of S-adenosylmethionine from L-methionine and ATP, is an ancient, highly conserved enzyme present in all three domains of life. Although the MAT enzymes of each domain are believed to share a common ancestor, the sequences of archaeal MATs show a high degree of divergence from the sequences of bacterial and eukaryotic MATs. However, the structural and functional consequences of this sequence divergence are not well understood. Here, we use structural bioinformatics analysis and ancestral sequence reconstruction to highlight the consequences of archaeal MAT divergence. We show that the dimer interface containing the active site, which would be expected to be well conserved across all three domains, diverged considerably between the bacterial/eukaryotic MATs and archaeal MATs. Furthermore, the characterization of reconstructed ancestral archaeal MATs showed that they probably had low substrate specificity which expanded during their evolutionary trajectory hinting towards the observation that all the modern day MAT enzymes from the three-kingdom probably originated from a common specific ancestor and then archaea MATs diverged in sequence, structure and substrate specificity. Altogether, our results show that the archaea MAT is an ideal system for studying an enzyme family which evolved to display high degrees of divergence at the sequence/structural levels and yet are capable of performing the same catalytic reactions as their orthologous counterparts.

## Introduction

Common descent is one of the fundamental aspects of Darwinian evolution. This theory emphasizes that modern day species diverged from a common ancestor ^*1*^. The same principle applies to enzymes: modern enzyme super-families across the three domains of life evolved from a set of enzymes that were already present in a last universal common ancestor (LUCA) dated over 3.5 billion years ago ^*2, 3*^. Many efforts have been made to infer the minimal set of LUCA enzymes ^*3, 4*^. The main hypothesis is that the evolutionary trajectory of enzymes (i.e. gene trees) would be closely associated with and influenced by the evolution of their respective host organisms (i.e. species trees) ^*5*^. However, in reality, gene evolution is much more complicated and disagreement between species trees and gene trees (non-congruence) can occur due to a wide range of factors such as gene duplication, Lateral Gene Transfer (LGT) ^*6, 7*^, and hybridization ^*8*^. Within this framework, a few studies have reported an unusual distribution of some enzymes among the three domains of life, whereby the archaeal enzyme shows divergence from more closely related eukaryotic and bacterial orthologs ^*9-12*^. One example is the Methionine AdenosylTransferase (MAT) enzyme. The unusual sequence similarity of MAT enzymes among the three kingdoms is striking; the archaeal enzyme is almost equidistant from the bacterial and eukaryotic enzymes with ∼20% sequence homology, respectively, while the bacterial and eukaryotic enzymes share greater than ∼60% homology ^*13*^.

MAT is a ubiquitous enzyme present across all three kingdoms of life that catalyzes the biosynthesis of S-adenosyl methionine (SAM) from L-Methionine and ATP. SAM is considered a pivotal methyl donor in nature right next to tetrahydrofolate derivatives (THF) ^*14*^. MAT is considered to be an essential enzyme and is required for cell growth ^*15*^. As many other enzymes in nature MAT is active as an oligomer (either as homo-oligomer or hetero-oligomer; Fig 1) ^*16*^. The oligomeric state provides a clear advantage over the monomeric state in terms of fitness, higher stability and functions ^*17, 18*^. The MAT homo-dimer is constituted by monomeric subunits (α-subunits) paired together along a large flat hydrophobic interface (henceforth referred to as large-interface - Fig 1) wherein the catalytic sites are enclosed ^*13*^. In case of a homo-tetramer, it exists as a dimer of homo-dimers, and so it has been suggested that the preferred functional quaternary structure is an oligomer (possibly an obligate dimer) ^*19*^.

**Figure 1.**
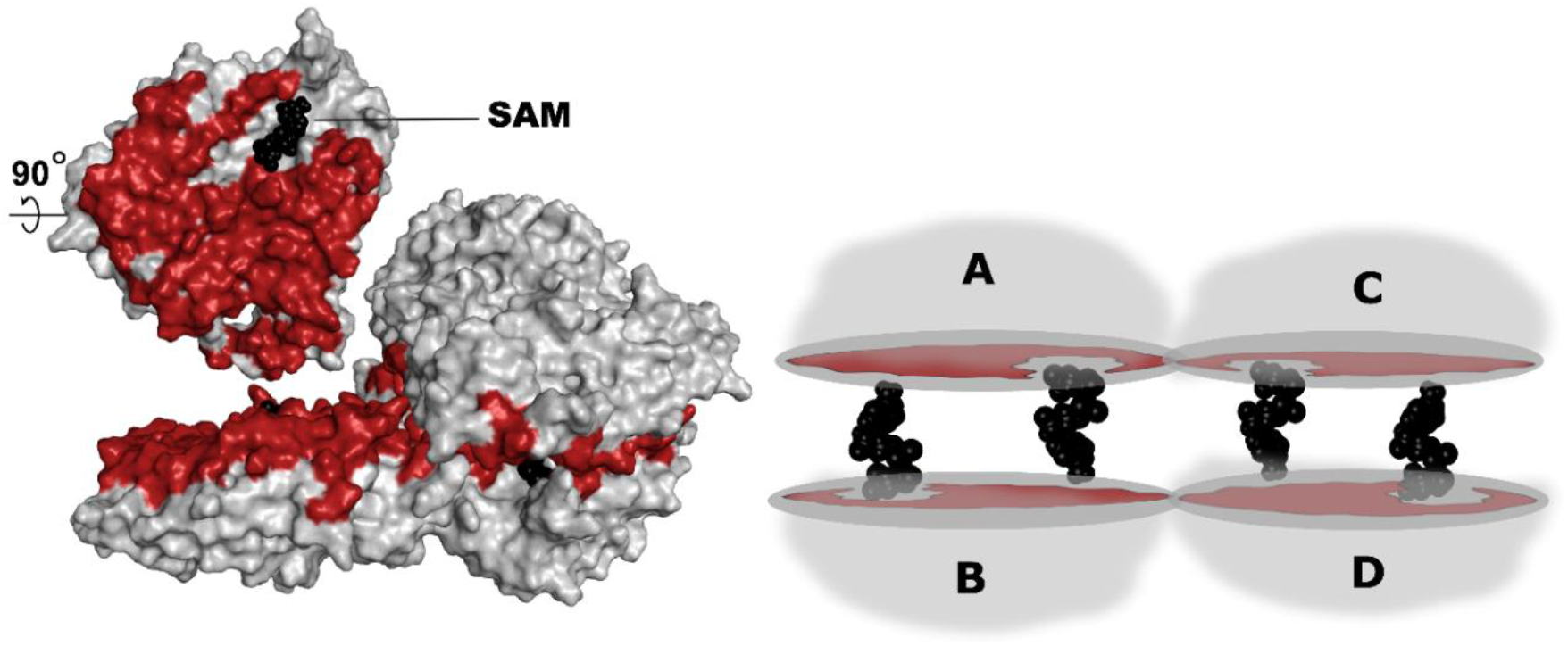
The MAT homo-tetramer configuration. Schematic representation of MAT homo-tetramer (right) and surface/cartoon view (left) – the large interface region (between chains A and B) is highlighted in red while SAM is highlighted in black. The large interface formed by chains A and B (also chains C and D) which has an area of ∼1900-2000 Å2 is constituted by ∼70-100 residues. Representative structures are utilized here, the archaea pfMAT structure (PDB: 6S83 from P. furiosus) utilizes 83 residues to give rise to the large interface and with 90 residues in case of bacteria eMAT (E. coli, PDB ID: 1RG9) and 77 in case of eukarya hMAT2A (H. sapiens, PDB ID: 4NDN).

It has been suggested that all extant MAT enzymes share a common ancestry ^*19*^ despite expressing a highly diverged form in archaea as mentioned above. This further indicates that adaptation of enzymes in course of evolution can lead to accumulation of mutations at the surface or interface regions especially near the catalytic sites. This phenomenon can allow for divergence and acceptance of a different substrate or function ^*20-25*^. Despite this contrast, it has been also established that archaea MAT can still perform the same catalytic reaction as their orthologous counterparts (from bacteria and eukarya) and so, the structural and functional consequences of this divergence are not well understood. These observations prompted us to take a closer look at the evolutionary trends adapted by the catalytic residues as well the large interface region in MAT. Therefore, we probed the putative evolutionary trajectory of archaea MAT in a more systematic manner by conducting investigations at various levels including sequence studies and structural comparisons, physio-chemical properties as well as performing ancestral sequence reconstruction (ASR). We further elucidate this potential evolutionary trajectory by resurrecting the ancestral archaea MAT sequences in the lab and characterizing their biochemical properties. In conclusion, we show that the archaea MAT is an ideal system for studying enzymes with high degrees of divergence at the sequence/structural level and yet perform the same catalytic reactions as their orthologous counterparts.

## Results and Discussion

### Probing the MAT sequence space

Herein, we conducted extensive searches to systematically probe the MAT sequence space across the three kingdoms. The sequences were collected by combining the outcomes of different databases (nr db, eggnog, orthodb, etc.). Upon compiling the first input dataset (∼ 800 sequences), we built sequence similarity networks (SSN). SSNs allow us to check MAT sequence space distribution across the three kingdoms (Fig 2B). During this analysis we considered that the archaea kingdom has not been sequenced as extensively as the other two kingdoms of life. Since phylogenetic trees may not be an optimum tool to highlight the overall sequence distribution pattern, we utilized two approaches - with EFI and CLANS ^*26*^ to visualize the sequence distribution pattern (S3 Fig). Despite the variation in the percentage sequence identity clustering ranging from 100-40% ID, the overall topology remains consistent with two major clusters representing: a) “bacteria + eukarya” and b) archaea. The SSN outcomes reflect the sequence identities as observed in nature, as archaea are nearly equidistant from the eukarya and bacteria at ∼20% sequence identity. SSN method provides an overall view of the topology for the dataset of interest and underline the uniqueness of the archaea kingdom.

**Figure 2.**
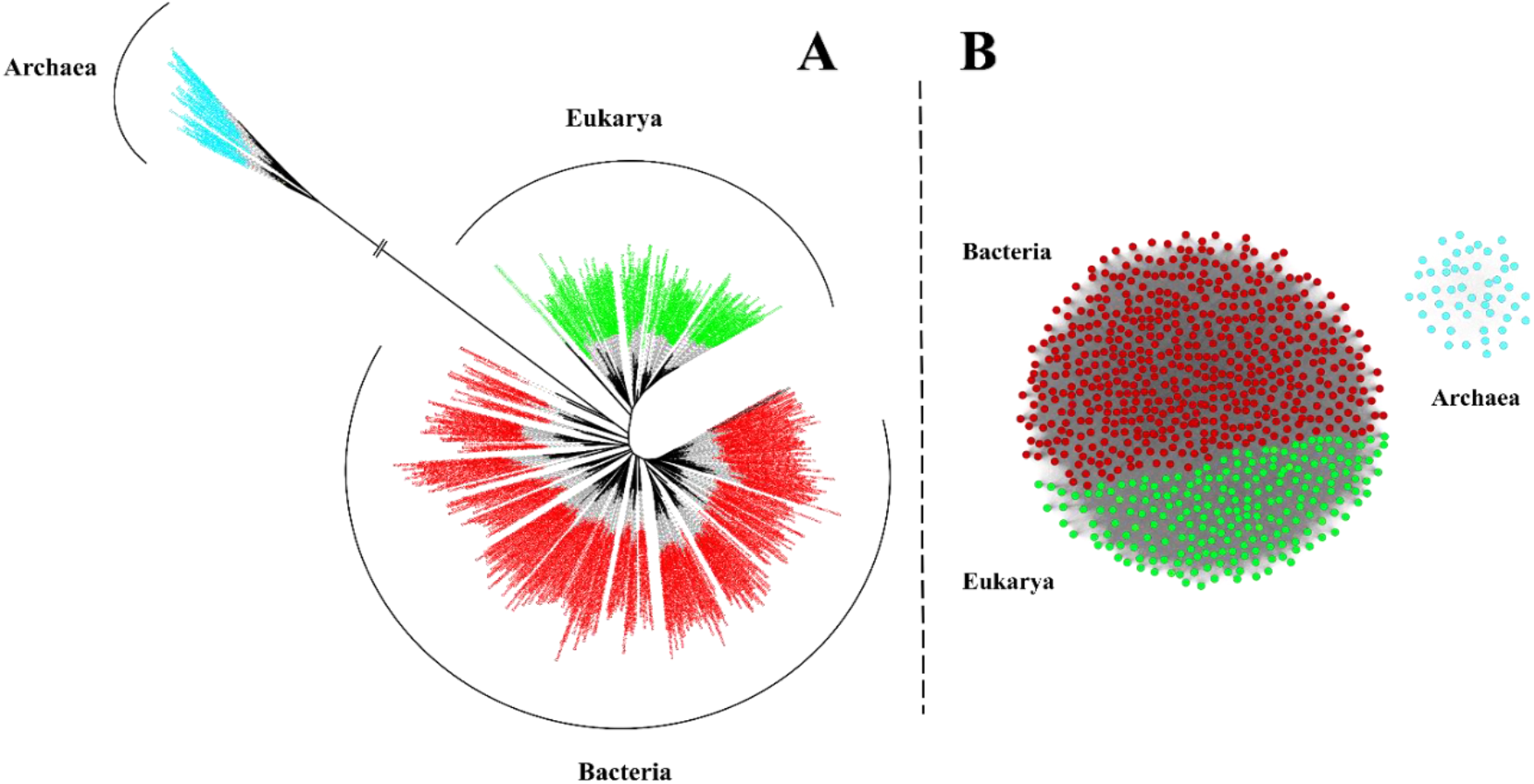
Schematic representation of observed sequence space in MAT. **(A)**. Unrooted Maximum Likelihood (ML) tree for the MAT from three kingdoms of life. This unrooted tree was built by implementing the ML method with IQ-Tree computer program and visualized using Figtree viewer program and iTOL server. The archaea MAT branches off thereby highlighting their divergence and this observation is further substantiated through sequence similarity network. The archaea MAT are represented in cyan, eukarya in green and bacteria in red. **(B)**. Sequence similarity network of MATs from the three kingdoms - sequences from eukarya (green) and bacteria (red) which form a distinct cluster while, the sequences from archaea (cyan) form a separate cluster. This SSN was created with the EFI-enzyme similarity server and visualization produced through cytoscape program. Clustering pattern within the network at different levels of sequence similarity (similarity %) can be observed in the supplementary information (S3 Fig).

### Inferring the ancestral MAT sequences

We reconstructed the phylogenetic relationships by utilizing MAT sequences from the three kingdoms of life (Fig 2A). In case of archaea, the two major phyla (crenarchaea and euryarchaea) were tested for the overall tree topology based on the MAT sequences (gene tree) as well as the corresponding 16s rRNA sequences (species tree) from the SILVA database (not shown here). Herein, the two topologies clearly indicate that the tree bifurcates into two distinct clades for crenarchaea and euryarchaea i.e. the major phyla are well classified into their respective monophyletic groups. Subsequently, we prepared another set of trees by utilizing bacterial MAT sequences as an outgroup in order to extract the putative common archaea ancestor (ArchAnc) (Fig 6). We also extracted the ancestral sequence for the crenarchaea (CrenAnc) and the euryarchaea (EuryAnc) at the designated nodes in the figure. The sequence similarities of the ancestral sequences (including interface residues and catalytic site residue comparisons) are detailed in other sections.

**Figure 6.**
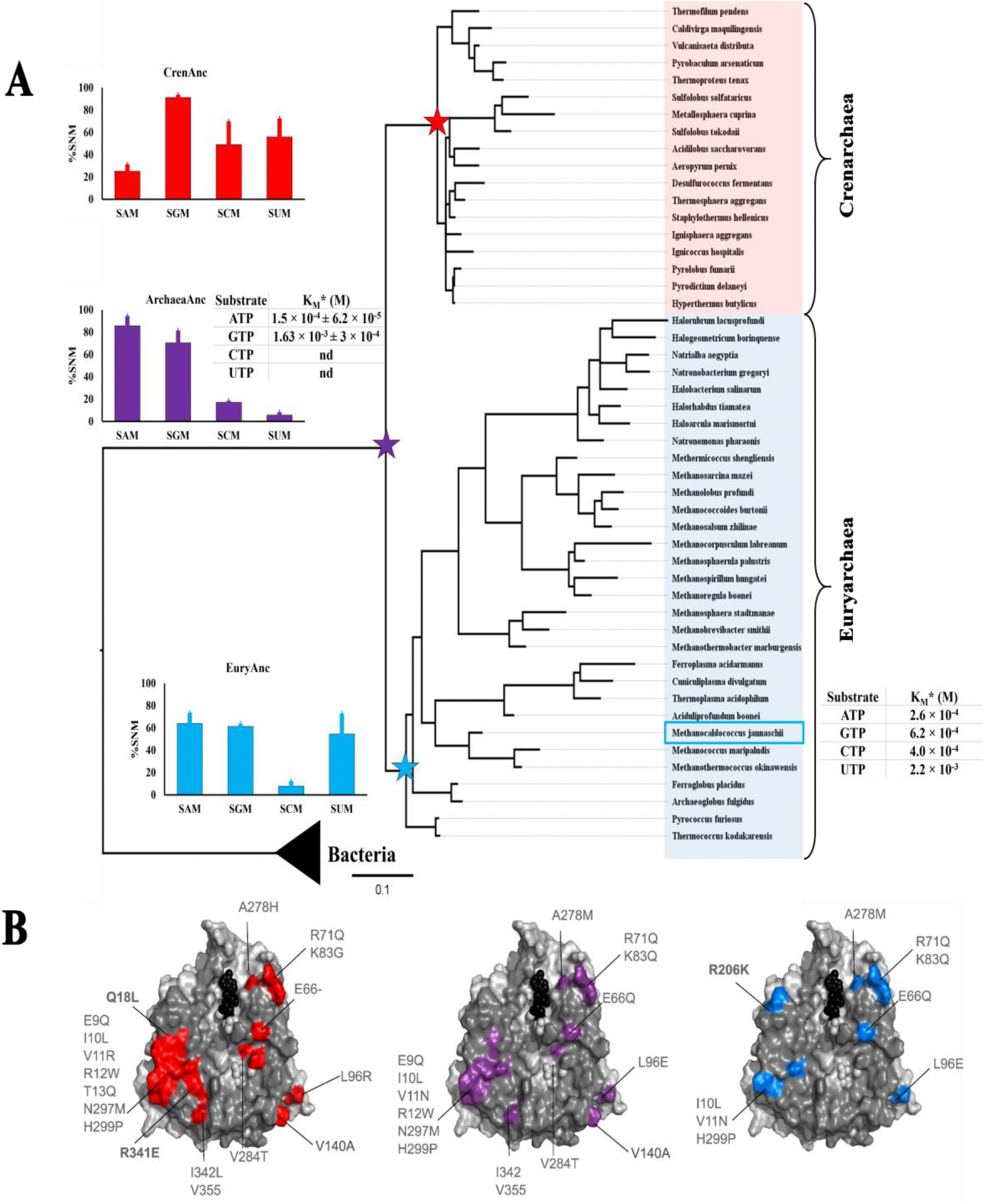
Phylogenetic tree of archaea MAT with bacteria and activity assays of the inferred ancestors. **(A)** The Maximum Likelihood phylogenetic tree topology of archaea MAT with bifurcation into two major clades i.e. crenarchaea and euryarchaea which are highlighted – bacteria was included as outgroup. This topology is further supported by another phylogenetic tree based on the 16s rRNA sequences from the corresponding species. Here, we highlight the activity of mjMAT as representative of kingdom archaea, with the data as reported previously by Lu et al, 2002. Ancestral sequences were inferred for the ancestor of Crenarchaea MAT (CrenArc in red star), Euryarchaea MAT (EuryArc in light blue star) and the common ancestor for the archaea MAT (ArcheaArc in violet star). End point assay for ancestral sequences activities were performed using corresponding enzyme (0.5 µM), NTPs (0.1-2 mM) and methionine (10 mM) in HEPES (100 mM) in HEPES (100 mM), KCl (50 mM), MgCl2 (10 mM), pH-8, at 37°C, 1h. The experiments were performed in duplicates. Tables showing the extrapolated K_M_ for ArcheaArc are reported (experimental condition: ArcheaArc 0.5 µM, ATP and GTP concentration in the range of 0.1 to 2 mM and methionine (10 mM) in presence of HEPES (100 mM), KCl (50 mM), MgCl2 (10 mM), pH 8 at 37°C. SAM and SGM production was analyzed by UPLC and data fitted to the Michaelis-Menten equation using GraphPad software. **(B)** Mutations highlighted on the interface region for the three ancestral MAT sequences in comparison to pfMAT (PDB: 6S83). CrenAnc MAT interface mutations highlighted in red, EuryAnc in blue and ArchaeaAnc in violet.

In the case of bacteria and eukarya MAT sequences, we prepared a separate phylogenetic tree (based only on the MAT sequences). Here, MAT sequences from these two kingdoms constitute two distinct clusters (also supported by PCA clustering) and this is also observed in the tree topology (Fig 3) and furthermore, this topology was utilized to infer the common MAT ancestors for eukarya (EukaAnc) and bacteria (BactAnc).

**Figure 3.**
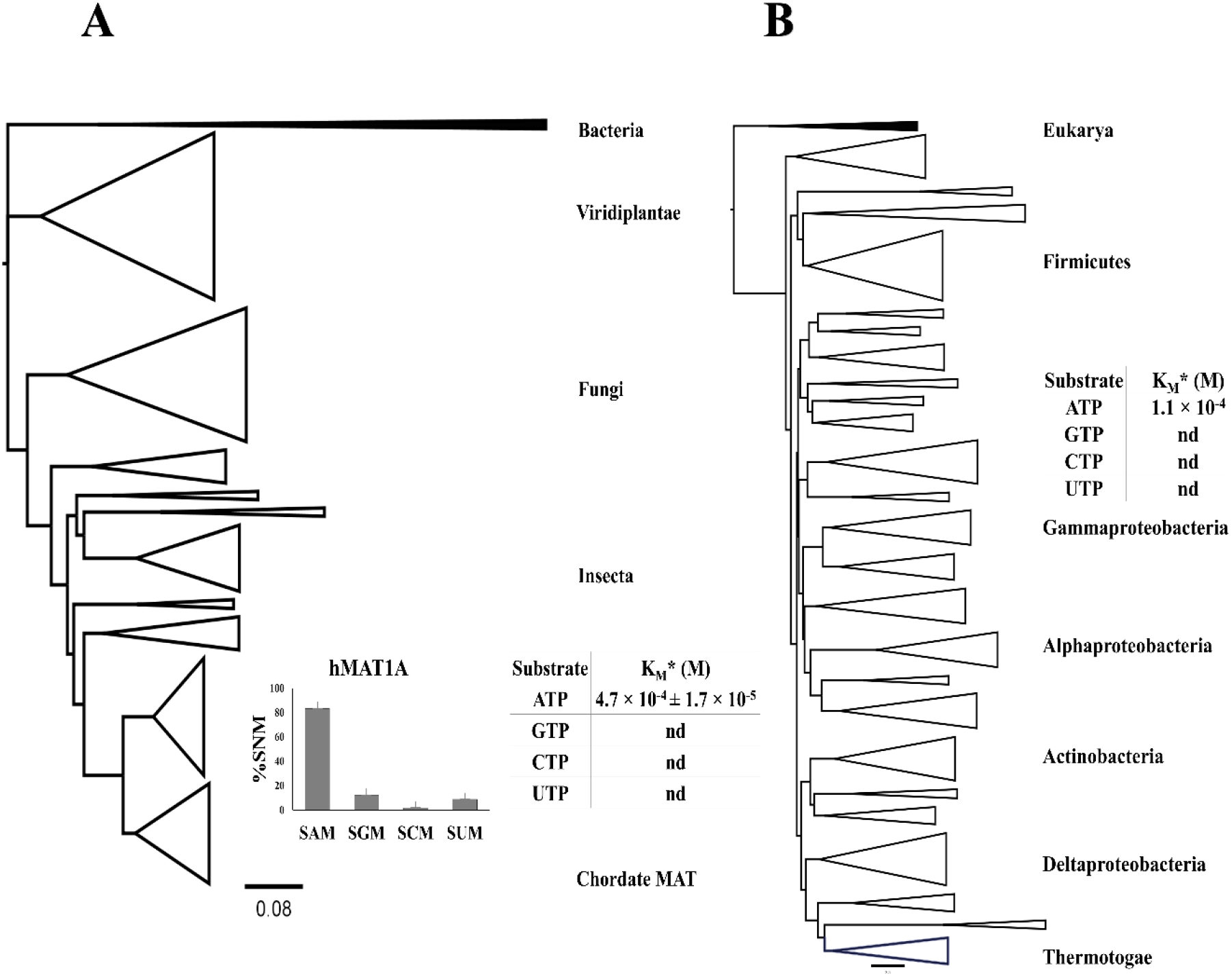
Phylogenetic trees based on the MAT sequences. **(A)**. Maximum/ likelihood phylogenetic tree of MAT sequences from kingdom eukarya with bacterial MAT sequences as an outgroup (highlighted in black). The major clades have been annotated according to their distribution in the tree topology. We reported the activity of hMAT1A as representative of kingdom eukarya. End point assay for hMAT1A activities were performed using hMAT1A (0.5 µM), NTPs (0.1-2 mM) and methionine (10 mM) in HEPES (100 mM), KCl (50 mM), MgCl2 (10 mM), pH-8, at 37°C, 1h. Experiments were performed in duplicates. Michaelis–Menten curves with detectable activity for ATP are reported in the Supporting information (S8 Fig). K_M_ for ATP was extrapolated and reported in tabular format. **(B)**. Maximum likelihood phylogenetic tree of MAT sequences from kingdom bacteria with eukarya MAT sequences utilized as an outgroup (highlighted in black). We also highlight the K_M_ of ATP from eMAT as a representative of kingdom bacteria as reported previously (Lu et al., 2002). These results suggest that both the enzymes hMAT1A and eMAT are not promiscuous as they display specificity towards ATP. The aforementioned trees were built by implementing the ML method with the IQ-tree program. Tree topologies were visualized with Figtree program.

### Extant MAT sequences vs ancestors: catalytic sites and evolutionary rates

Herein, we probe the evolutionary trend(s) for amino acid sites in the MAT enzymes across the three kingdoms of life. These trends were visualized by mapping the interface sites as well as the catalytic sites on a representative X-ray structure (Fig 4, S4 Fig). Additionally, we also probe the level of conservation for catalytic site residues in extant as well as ancestral sequences.

**Figure 4.**
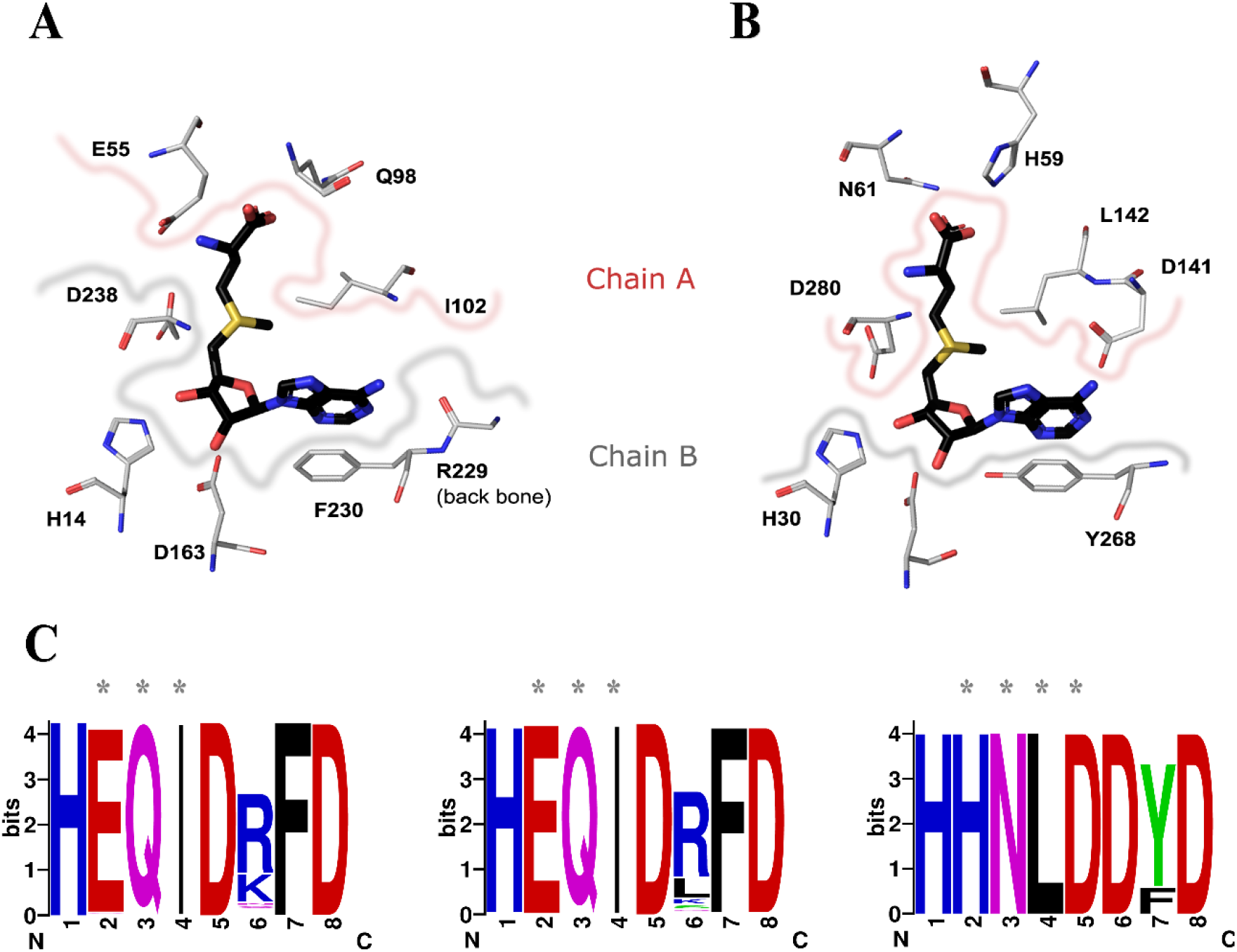
Catalytic residues involved in the interaction with SAM. **(A)**. Here the S-Adenosyl Methionine (SAM) is highlighted in black while the active site residues from eMAT (PDB: 1RG9) are highlighted in metal grey. The residues derived from chain A are demarcated with a pinkish border while the corresponding residues from chain B are demarcated with a greyish border. Similarly, the residues from pfMAT (PDB: 6S83) interacting with SAM have also been highlighted in **(B)**. **(C)**. The corresponding catalytic residues (based on structural alignment) from all three kingdoms of life are highlighted as web logos (https://weblogo.berkeley.edu/logo.cgi) and the residues derived from the second monomeric subunit are marked by ‘*’

#### Catalytic sites (active site residues)

in the case of archaea, we observe that the MAT catalytic site residues are very well conserved among the two major branches i.e. crenarchaea and euryarchaea (residue numbers correspond to pfMAT structure: PDB:6S83, (17)). These include the following residues involved in interaction with SAM from chain A: H59, N61, D141, L142, D280 and from chain B: H30, Y268. Furthermore, these catalytic site residues are completely conserved in the putative archaea ancestor. A similar trend can also be observed in the case of both eukarya and bacteria. This provides us with a key insight that despite observing differences in the catalytic site residues, MAT enzymes from the three kingdoms can still essentially perform the same catalytic reaction (Fig 4).

#### Evolutionary rates

Besides the conservation level of the catalytic site residues, we also probe the evolutionary rate distribution of all residues in the MAT structure (S4 Fig). Interestingly, most sites located along the large interface tend to be slowly evolving (highlighted in blue) while the situation changes as we move further away from the large interface. For instance, some of the residues located along the surface α-helices and loop regions tend to experience higher evolutionary rates. This pattern can be seen in MAT enzymes from all three kingdoms of life – thereby indicating that the conservation at large interface is probably critical for enzyme function and so we do not find evidence to show that the archaea MAT interface residues retain high evolutionary rates. The stabilization of the homo-oligomer state most likely plays a key role in the evolutionary dynamics of this interface.

### Large interface: charge and hydrophobicity distribution

In case of MAT enzymes, the constituent monomers pair up in an inverted configuration by exposing the α-helices towards the surface and the β-strands interact to form a hydrophobic isologous large interface that harbor the catalytic residues – thereby making this homo-dimer the obligate functional unit ^*27*^. Herein, we notice that divergence is not just limited to catalytic residues, or the residues located close to the substrate, but most of the inter-dimer interface (large-interface) residues are also quite diverged across the three domains of life, for instance - in case of bacteria (eMAT, PDB: 1RG9, ^*28*^) and eukarya (hMAT1A, PDB:6SW5, ^*29*^) - 38 out of 51 aligned residues are identical. In contrast archaea (pfMAT, PDB: 6S83) shares only 12 identical residues each with bacteria (eMAT) and eukarya (hMAT1A) (S1 Fig, S2 Fig). This further prompted us to perform a systematic analysis of the large interface region across the three domains of life and so, we conducted an analysis of the charge and hydrophobicity distribution across the large interface and also compare them with the ancestral sequences to gain more insights into evolutionary patterns. We proceed by studying 17 experimentally solved MAT structures from Protein Data Bank to probe the physio-chemical properties by creating two datasets: i). identifying 51 structurally aligned positions along the large interface (which includes up to five non-interface residues as well) and ii). Identifying 24 structurally aligned ‘interface residues only’ positions.

#### Large interface region (extant)

Based on the aforementioned 51 structurally aligned positions, it is clear that even at a sequence level the MAT large interface of bacteria and eukarya has a striking sequence identity of ∼50-70%. However, the archaea interface sequences are nearly equidistant from the other two kingdoms at ∼15-30% sequence identity, respectively, despite having a comparable size in terms of area (∼1660-2970 Å2). The hierarchical clustering plots (based on pvclust package in R, Fig 5, ^*30*^) show that the sequence identities further dictate physio-chemical properties as well. For instance, a look at both the P-values provided by pvclust i.e. au (approximately unbiased) p-value and bp (bootstrap probability) value for comparison of the clusters reveals that the hydrophobicity and charge distribution are cluster together for bacteria and eukarya MAT structures with archaea MAT structures clustering off separately. An additional analysis was also conducted for the second dataset with an alignment of 24 ‘interface-only’ residues (S5 Fig) and in this case too, we observe similar results as archaea MAT structures cluster off separately.

**Figure 5.**
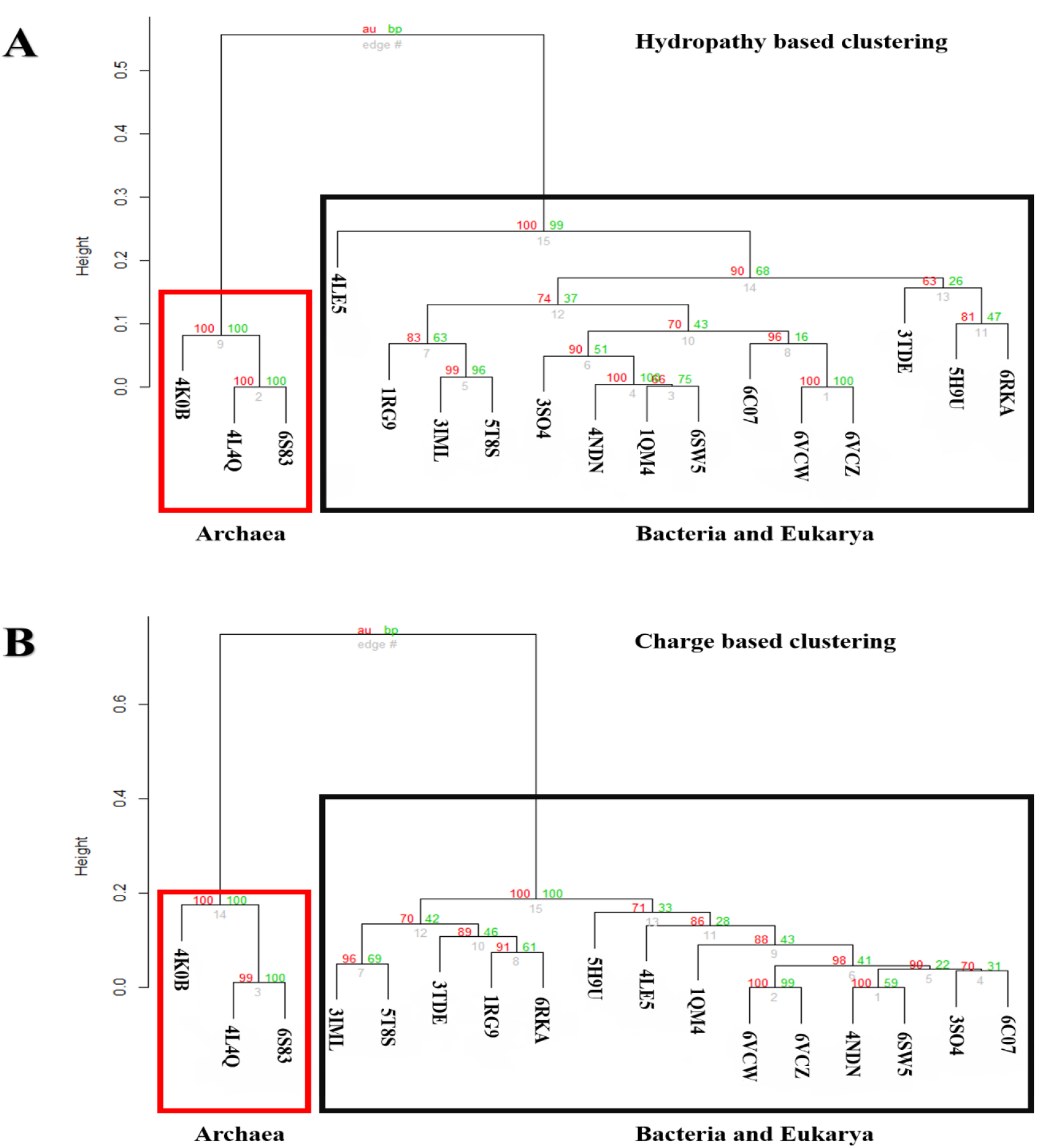
Clustering analysis of hydropathy **(A)** and charge **(B)** of representative structures from the three kingdoms of life. Clustering analysis of the physiochemical properties (Hydropathy and Charge) of 51 structurally aligned large-interface residues from chain A of 17 MAT structures (PDBs list in the SI). Based on the 51 aligned positions we calculated the per-site hydropathy with the help of Kyte-Doolittle scale from the protscale server and per-site charges were calculated with EMBOSS charge server for the aforementioned 17 MAT structures. Subsequently, we conducted clustering analysis with the help of pvclust package in R. This further provides a statistical score in terms of AU (approximately unbiased scores depicted in red) p-value and BP (bootstrap probability scores depicted in green) value for comparison of the clusters which reveals that the hydropathy **(A)** and charge distribution **(B)** cluster together for bacteria and eukarya MAT structures with archaea MAT structures clustering separately. In both the cases the archaea cluster is provided with high support values with both the AU and BP parameters, 100 % support for a distinct archaea cluster with respect to the two physiochemical properties each

#### Ancestral vs extant (large interface)

Herein, we made a comparison of the large interface residues extracted from the ancestral sequences against their extant counterparts. A comparison of the large interface residues from the bacterial MAT ancestor with the corresponding residues from eMAT highlights a difference of 13 residues out of 51 residues (S2 Fig). Likewise, we also observe similar changes for eukarya: hMAT1A vs eukarya MAT ancestor with 7 residue changes (out of 51 residues) and for archaea: pfMAT vs archaea MAT ancestor with 6 changes (out of 51 residues).

In conclusion, these observations provide a crucial insight that the sequence identities further influence the distribution of corresponding physio-chemical properties. Therefore, the hydrophobicity and charge distribution as described the hierarchical clustering (based on pvclust) are important in understanding the evolutionary trends adapted by the large interface. However, we do observe that the catalytic sites remain conserved to some extent across the MAT enzymes.

### Structural comparison supports sequence divergence

Structural similarity searches were carried out to explore the MAT structural neighborhood by implementing DALI-server based ^*31*^ searches against the PDB database. Representative MAT structures, from three kingdoms of life were utilized as templates to conduct searches. Interestingly, the output results suggest that the immediate structural neighborhood (∼2Å RMSD) is almost exclusively constituted by other MAT structures. This mirrors our observations in case of sequence-based searches as well (for each kingdom) i.e. the top-hits reported during the NCBI BLAST searches are constituted by MAT sequences. These observations could further point towards a shared common ancestral fold with a highly specialized and divergent form in archaea. Furthermore, since MAT is composed of three constituent domains – Domains I-III (also described as N-terminal domain, C-terminal domain and Central domain). DALI searches against the PDB database was carried out by utilizing individual domains as search queries and the resulting output is quite similar to the aforementioned DALI searches.

We further extended the comparison by utilizing the constituent MAT domains. The MAT representative structures were segregated into constituent domains by implementing the SCOPe defined boundaries (for the domains I-III) in case of eukarya and bacteria. The three constituent domains were then compared among themselves (within the kingdom) as well as across (among the kingdoms). Unfortunately, the PDB dataset for archaea MAT (PDB: 4L4Q, ^*32*^; 4K0B, ^*33*^; and 6S81) are currently not classified into constituent domains in SCOPe v2.07, therefore we performed this analysis for archaea MAT based on the demarcation as reported in literature (S6-S7 Fig). Similarly, this domain analysis was also conducted for MAT across the three kingdoms wherein the constituent domains were probed for one to one relationship to elucidate structural homology. In this case too, the domains were compared at a structural and sequence level against each other (S6-S7 Fig). These observations provide the insight that at least in case of bacteria and eukarya the constituent MAT domains display high level of homology with archaea expressing a highly diverged form as reflected in the sequence searches as well.

In conclusion, our structural comparison results support the observations made at a sequence level and indicate a common descent from a shared ancestral fold with high level of divergence in MAT domains within and across the three kingdoms of life.

### Change in substrate specificity along the archaea phylogenetic tree

It has been previously reported in literature that the archaea mjMAT is promiscuous towards different nucleotides i.e. it is able to accept different NTPs while the bacterial ortholog eMAT is specific for the double ring adenine nucleotide base ^*13, 34*^. This further prompted us to study the specificity for the purine bases (of ATP and GTP) within the archaea phylogenetic tree and see if this information could provide us with more clues towards the origin and evolution of MAT (Fig 6). We expressed and tested the activity of the ancestral archaea MAT sequences and compared them against the representative extant MAT sequences from the three kingdoms of life (Fig 3, Fig 6). Data about specificity of the representative bacteria and archaea extant MAT were already reported in literature ^*13, 34*^. As representative of the eukarya we choose hMAT1A. Since our purpose was to analyze the change in specificity within kingdom archaea - we expressed and tested the activity for the ancestral MAT sequences for euryarchaea (EuryAnc), crenarchaea (CrenAnc) and the common archaea ancestor (ArchaeaAnc) (S8-10 Fig, S13-15 Fig, S18 Fig). CrenAnc displays similar promiscuities towards all the bases much alike the extant MAT enzyme (Fig 6), while EuryAnc loses promiscuity towards the cytidine triphosphate (CTP). Interestingly, ArchaeaAnc acquired specificity for Adenine triphosphate and Guanidine triphosphate (ATP and GTP respectively). In contrast, eukarya hMAT1A retains specificity for the ATP substrate (S12 Fig, S17 Fig, S18 Fig). Unfortunately, the bacterial and eukaryotic ancestral MAT sequence could not be expressed and so, their data are missing here. Furthermore, in the archaea phylogenetic tree we detect a change in specificity, starting from a more specific ancestor towards a more promiscuous extant enzyme. We observe that the specificity for ATP vs GTP goes from a ten-fold difference in the ancestor ArchaeaAnc to three-fold difference in mjMAT, which indicates that the ancestor was specific towards purine bases but through the course of evolution modern day archaea MAT became more promiscuous towards different bases. Therefore, this data supports the observation that all the modern day MAT enzymes from the three-kingdom probably originated from a common specific ancestor and then archaea MATs diverged in sequence, structure and substrate specificity.

### Additional factors influencing the evolutionary trajectory of MAT

In this study, we have shown that various factors could have contributed towards the unusual evolutionary trajectory of the MAT enzyme across the three kingdoms of life. However, there are some additional aspects could have played a significant role in shaping the evolutionary trajectory of MAT enzyme in archaea, these could include: codon usage bias, tRNA bias, or an alternative/distinct SAM metabolism. It has been shown that in general, the archaea genomes display a higher GC-rich tendency in contrast to bacterial and eukaryotic genomes. This tendency has a direct influence in the codon usage. Codon usage bias has also a direct association with tRNAs abundance for protein translation optimization ^*35, 36*^. This includes avoiding slowly translated codons and utilizing codons with the most cognate abundant tRNAs from the genome. Additionally, in highly expressed genes, favored codons are easily recognized by the abundant tRNA molecules ^*37-39*^. This bias could be further augmented by the time and speed of gene expression as it could also play a critical role in the presence of more abundant and less diverse tRNAs as well ^*40*^. Another possible factor that might have influenced the divergence of archaea MAT is a different SAM metabolism. Although, we acknowledge the presence of a different methyl donor in archaea (tetrahydromethanopterin - THMPT) but no clear connection to the SAM metabolism was found to support the hypothesis of the alternative MAT evolutionary trajectory. Additionally, since their divergence from LUCA, the selection and adaptive forces operating on bacterial/eukaryotic and archaeal clades may have differed substantially resulting from a variety of factors including: different environmental and metabolic constraints, differences in of regulation of activity, changes in oligomerization state as well as intracellular turnover. Therefore, we anticipate that these additional factors could also play a key role in guiding the evolution of highly diverged archaea MAT as well as other enzymes.

## Conclusion

In closing, we expected the large interface region from MAT to be well conserved across the three domains of life primarily due to these main reasons: i) it accommodates the catalytic residues; ii). facilitates the same catalytic reaction and, iii) it is composed of a large flat hydrophobic region constituted by β-sheets which are known to evolve more slowly compared to α-helical regions - these are localized at the surface in the case of MAT ^*41, 42*^. However, our results clearly suggest that the pivotal large interface region is surprisingly diverged in case of archaea MAT in contrast to bacteria/eukaryotic MAT. We anticipate that future advances in the genome sequencing projects - especially related to kingdom archaea-could shed light on other enzymes that evolutionary behave similarly

## Methods

### Sequence data collection

MAT sequences from Methanocaldococcus jannaschii (uniport: **Q58868**, METE_METJA), Escherichia coli (uniprot: **Q58605**, METK_METJA), and MAT1A from Homo sapiens (uniprot: **Q00266**, METK1_HUMAN) were used for conducting NCBI BLAST ^*43*^ searches across the non-redundant database. Furthermore, the database search conditions were filtered based on NCBI recommended e-value cutoff i.e. 1e-5 and only search results with query coverage >90% and sequence identity >55% were considered in constructing the initial sequence dataset. The collected sequences were then subjected to reciprocal BLAST searches to confirm orthologous relationship(s) as it is a common computational method for predicting putative orthologues. METK sequences for archaea from the phylogenetic group *asgard* (superphylum) were not considered in this study owing to the lack of taxonomic classifications associated with the sequences that were reported as hits in the NCBI BLAST searches.

#### Sequence analyses

The CD-HIT program ^*44*^ was utilized to reduce the sequence redundancy by clustering at 80% sequence identity cutoff threshold with default settings. The Gblocks program ^*45*^ was implemented to identify highly conserved sites across sequence alignments with assistance from the secondary structure information (from experimentally solved crystal structures) to guide the alignments. Sequence alignment was carried out by using the MAFFT computer program ^*46*^ followed by manual curation to check for any errors.

### Phylogenetic analysis and ancestral sequence reconstruction

Modeltest computer program ^*47*^ was implemented to pick out the best evolutionary model for sequence alignments based on BIC (the Bayesian Inference Criterion). The LG model ^*48*^ with invariant sites (+I) for discrete gamma categories, (+G4) was selected to construct ML phylogenetic trees. Initially, we constructed Maximum Likelihood (ML)-based trees with the IQ-TREE program ^*49*^ (using 10,000 bootstrap replicates) for the datasets to inspect the topology distribution within the dataset. Confidence level for the nodes was assessed with Felsenstein’s bootstrap method ^*50*^ and the consensus tree was redrawn using Figtree (http://tree.bio.ed.ac.uk/software/figtree/). Tree file from IQ-tree program output was parsed to obtain the ancestral sequences at highlighted nodes. Also, phylogenetic program MEGA-X was consulted to check for the ancestral sequences ^*51*^. The NCBI taxonomy database was also consulted to check for the major phylogenetic classifications^*52*^.

### Evolutionary rates calculations

The site-specific evolutionary rates were inferred through the IQ-TREE program by implementing the estimated model parameters and further applying an empirical Bayesian approach to assign site-rates as mean over rate categories.

### Hierarchical clustering with pvclust

An R language package was utilized for assessing the hierarchical clustering pattern as it provides statistical support for each cluster through P-values. It provides two types of p-values: AU (Approximately Unbiased) p-value and BP (Bootstrap Probability) value. AU p-value, which is computed by multiscale bootstrap resampling, is a better approximation to unbiased p-value than BP value computed by normal bootstrap resampling.

### Molecular modeling and structural studies

Ancestral sequences (of interest) were modeled by using the Swiss-model server at Expasy server (https://swissmodel.expasy.org/). 3D models were visualized with Pymol (https://pymol.org/2) to identify, tabulate and study the key interaction residues. The overall quality of the 3D models depends directly on the shared sequence identity between the target sequence and the sequence of the template structure, therefore we utilized template structures with high sequence similarity. Interface residues were identified through Pymol and PISA server (https://www.ebi.ac.uk/pdbe/pisa/). Structural superposition studies were carried out with mtm-align server ^*53*^.

### Charge and hydrophobicity calculations

Per-site hydrophobicity and hydrophilicity calculations were carried out by using the ProtScale program ^*54*^ from the Expasy server (https://web.expasy.org). Here, we implemented the default “Kyte and Doolittle” scale ^*55*^ for the calculations with residue window size 3, while the rest of the settings were left as default. Charge distribution per site were calculated using the charge server from EMBOSS (http://www.bioinformatics.nl/cgi-bin/emboss/charge). Here, we implemented a window length of size 3 and the rest of the settings were left as default.

#### Sequence similarity network (SSN) studies

Sequence similarity networks were constructed based on the output obtained from the EFI-EST (Enzyme Similarity Tool) server ^*56*^ and subsequently the cytoscape program (https://cytoscape.org/) was utilized to explore the SSNs via the organic layout. Here, we managed the initial dataset by CD-HIT based clustering by selecting a sequence identity threshold of 80 percent. The “organic” layout algorithm, was chosen for graphical clustering.

### Experimental methods

ATP, GTP, CTP, UTP, methionine, S-adenosylmethionine (SAM), HEPES, MgCl2, KCl, isopropyl-1-thio-β-D-galactopyranoside (IPTG), Tris HCl, Na2HPO4, NaH2PO4, potassium phosphate, NaCl, imidazole, β-mercaptoethanol, dithiothreitol (DTT), kanamycin, glycerol, NaOH, HCl, ammonium acetate, bacto agar, bacto tryptone, bacto yeast extract all other chemicals and HPLC grade solvents were purchased from commercial sources and used as supplied unless otherwise mentioned. Page Ruler prestained protein ladder, 10 to 180 kDa was purchased from Thermo Fischer scientific. BL21 (DE3) competent cells were purchased from New England biolabs (NEB). Benzonase, complete His-Tag Purification Resin (NiNTA) were purchased from Sigma Aldrich. Protein inhibitor cocktail (PIC) and Lysozyme were purchased from Nacalai Tesque, INC. 12% Mini-PROTEAN TGX Precast Protein Gels, 12-well from BIO-RAD. Amicon centrifugal filters were purchased from Merck. In-Fusion HD cloning kit was purchased from Takara. All the experiments were performed using ultrapure water purification system from a MilliQ Integral MT10 type 1 (Millipore).

#### Reaction Conditions for the formation of SNM analogs

ATP/GTP/CTP/UTP (5 mM), methionine (10 mM), HEPES (100 mM), MgCl2 (10 mM), KCl (50 mM) and MAT (20 µM) were mixed in water, pH was adjusted to 8 with 10% NaOH. The reactions were incubated at 37° C for eukarya and 55 °C for archaea in thermomixer comfort (Eppendorf) for 1 hr. Reaction was quench by acetonitrile followed by centrifugation at 12,000 RPM for 5 min to precipitate the enzymes. Finally, supernatant was filtered through 0.22 µm filter (Merck) and injected in UPLC for analysis (Waters UPLC Acquity H class).

#### UPLC analysis for SNM analog detection

Diluted reaction aliquots were analyzed by UPLC (Waters UPLC Acquity H class) using a HILIC column (SeQuant ZIC-cHILIC 3 µm,100 Å 150 x 2.1 mm PEEK coated HPLC column). An isocratic method was used with solvent A (100 mM ammonium acetate, pH 5.3) 35% and solvent B (acetonitrile) 65% for 15 min. Each injection was 3 µL with a flow rate of 0.2 mL/min and detected at 260 nm.

## Accession codes

MAT sequences from Methanocaldococcus jannaschii (uniport: **Q58868**, METE_METJA), Escherichia coli (uniprot: **Q58605**, METK_METJA), and MAT1A from Homo sapiens (uniprot: **Q00266**, METK1_HUMAN)

## Acknowledgments

Financial support by the Okinawa Institute of Science and Technology is gratefully acknowledged. We also thank Dr. Benjamin Clifton and Professor Mark S. Johnson for a critical reading of the manuscript and providing feedback to improve the quality of the manuscript.

## Author contributions

BPSC and PL conceived the idea of the study. BPSC, MHG, and DM performed the experiments. BPSC, MG, STP, and PL analyzed the data. BPSC, MHG, STP and PL wrote the manuscript and prepared the figures.

## Data availability statement

The data underlying this article are available in the article and in its online supplementary material. Also, any other data underlying this article will be readily shared on request to the corresponding author.

## Abbreviations

(LUCA): Last Universal Common Ancestor
(LGT): Lateral Gene Transfer
(MAT): Methionine AdenosylTransferase
(SAM): S-adenosyl methionine
(ATP): Adenosine triphosphate
(GTP): Guanosine triphosphate
(CTP): Cytosine triphosphate
(UTP): Uridine triphosphate
(ASR): Ancestral Sequence Reconstruction
(SSN): Sequence Similarity Networks
(PCA): Principal component analysis

## For table of Contents Only

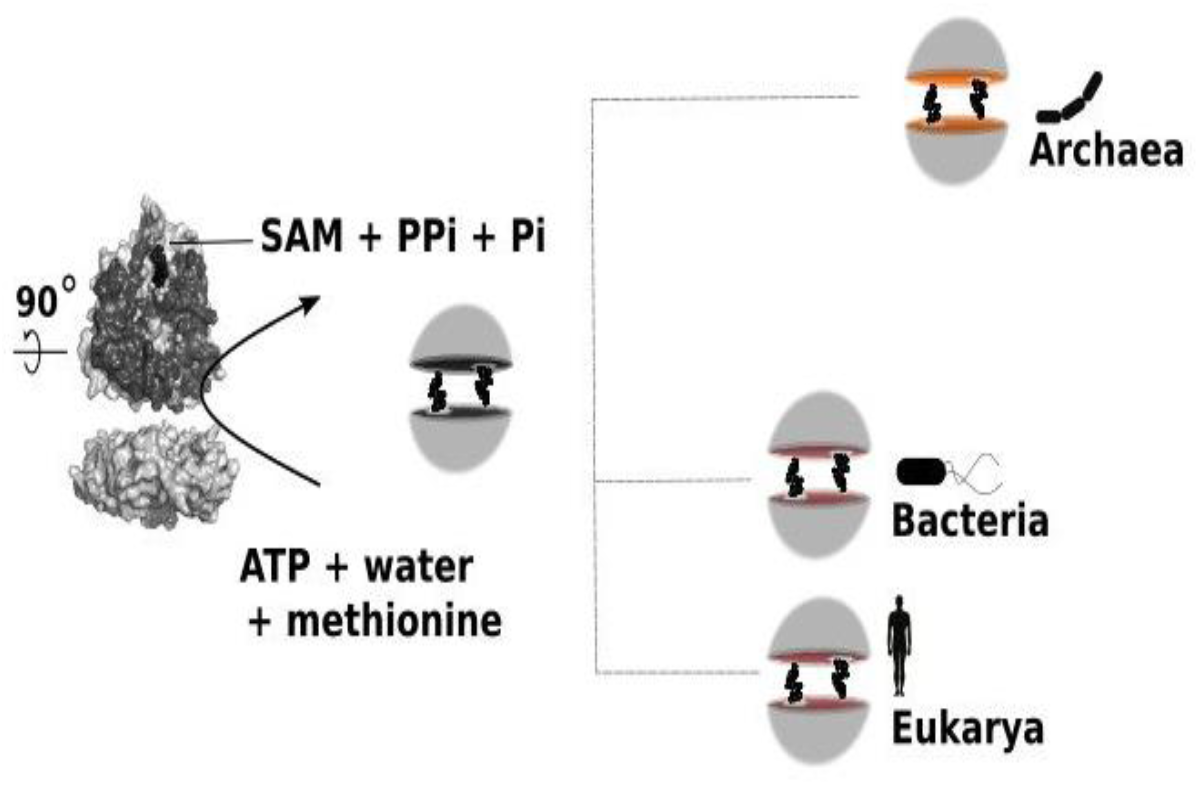

