## supplementary information for "Consequences of the divergence of Methionine AdenosylTransferase"

**Contents:**

**Supplementary information text**

**Supplementary figures**

**Supplementary references**

**Supplementary information text**

**Primers for infusion cloning**

| Name | Froward Primer | Reverse Primer |
|---|---|---|
| Pet28a_linearization | 5'-TCCGTCGACAAGCTTGCGGCCGCAC-3' | 5'-GCGGCACCAGGCCGCTGCTGTGATG-3' |
| MAT_ amplification | 5'-GCAAGCTTGTCGACGGAGCTCGAATTC-3' | 5'-GCAAGCTTGTCGACGGAGCTCGAATTC-3' |

Sequences of the enzymes used in this study

**hMAT1A sequence**

MNGPVDGLCDHSLSEGVFMFTSESVGEGHPDKICDQISDAVLDAHLKQDPNAKVACETVCKTGMV
LLCGEITSMAMVDYQRVVRDTIKHIGYDDSAKGFDFKTCNVLVALEQQSPDIAQCVHLDRNEEDV
GAGDQGLMFGYATDETEECMPLTIILAHKLNARMADLRRSGLLPWLRPDSKTQVTVQYMQDNGAV
IPVRIHTIVISVQHNEDITLEEMRRALKEQVIRAVVPAKYLDEDTVYHLQPSGRFVIGGPQGDAG
VTGRKIIVDTYGGWGAHGGGAFSGKDYTKVDRSAAYAARWVAKSLVKAGLCRRVLVQVSYAIGVA
EPLSISIFTYGTSQKTERELLDVVHKNFDLRPGVIVRDLDLKKPIYQKTACYGHFGRSEFPWEVP
RKLVF

**MjMAT sequence**

MRNIIVKKLDVEPIEERPTEIVERKGLGHPDSICDGIAESVSRALCKMYMEKFGTILHHNTDQVE
LVGGHAYPKFGGGVMVSPIYILLSGRATMEILDKEKNEVIKLPVGTTAVKAAKEYLKKVLRNVDV
DKDVIIDCRIGQGSMDLVDVFERQKNEVPLANDTSFGVGYAPLSTTERLVLETERFLNSDELKNE
IPAVGEDIKVMGLREGKKITLTIAMAVVDRYVKNIEEYKEVIEKVRKKVEDLAKKIADGYEVEIH
INTADDYERESVYLTVTGTSAEMGDDGSVGRGNRVNGLITPFRPMSMEAASGKNPVNHVGKIYNI
LANLIANDIAKLEGVKECYVRILSQIGKPINEPKALDIEIITEDSYDIKDIEPKAKEIANKWLDN
IMEVQKMIVEGKVTTF

**ArchaeaAnc sequence**

MRNIVVEQLNWTPVEEQQVELVERKGLGHPDYIADGISEAVSRALCKYYLERFGTILHHNTDQVQ
VVGGQASPRFGGGEVIQPIYILLSGRATTEVDGEKVPIGTIALKAAKDWLRENFRFLDPERHVII
DCRIGQGSADLVGVFERGKSVPLANDTSFGVGFAPLSTTERLVFETERFLNSKFKKKYPAVGEDI
KVMGLRRGKKITLTIAAAMVSRFVKDMDEYLSVKEEVKDAVQDLASKYTPYDVEVYVNTADKPEK
GIFYLTVTGTSAEMGDDGSTGRGNRCNGLITPMRPMSMEATAGKNPVSHVGKIYNILANQIAQRI
YEEVKGVKEVYVRLLSQIGKPIDQPLIANVQVIPEDGYLTSDMKREIEAIADEWLANITKITEMI
LEGKVSLF

**EuryAnc sequence**

MRNIVVEELNRTPIEEQQVELVERKGIGHPDSIADGIAEAVSRALCKEYMERFGAILHHNTDQVQ
VVGGQAHPRFGGGEVIQPIYILLSGRATKEVDGEKIPVDTIALKAAKDYLRETFRHLDLERHVII
DCRIGQGSVDLVGVFNRQKPVPLANDTSFGVGYAPLSETERLVFETERFLNSEFKKKYPAVGEDI
KVMGLRKGDKITLTIAAAMVDRYVSNMDEYLEVKEEIKDAVKDLASKYTDREVEVYVNTADDPEK
GCFYLTVTGTSAEMGDDGSVGRGNRCNGLITPNRPMSMEATAGKNPVSHVGKIYNILANQIAQDI
AEEVEGVKEVYVRILSQIGKPIDQPLVASVQVIPEDGYSISDMEREVKEIADEWLANITKITEMI
LEGKISVF

**CrenAnc sequence**

```
MRNIVVEQLRWQPVEELQVELVERKGLGHPDYIADAISEAASRELSKYYLERFGTILHHNLDKVL
VVGGQASPRFGGGEVIQPIYILVSGRATTEVDGEKVPIGTIILKAAKDWIRENFRFLDPERHVII
DYRVGQGSADLVGIFELGKSVPLANDTSFGVGFAPLSTTERLVFETERLLNSKFKAKFPAVGEDV
KVMGLRRGKKIKLTIAAAIISRFVKDMDEYLSVKEEVKDAVLDLASKIAPYDVEVYVNTADKPEK
GIFYLTVTGTSAEHGDDGATGRGNRANGLITPMRPMSMEATAGKNPVSHVGKIYNVLANQIAQRI
YEEVKGVKEVYVELLSQIGKPINEPLIANVQVIPEEGELTSDMKREIEAIADEELDRITKITEMI
LEGKVSLF
```

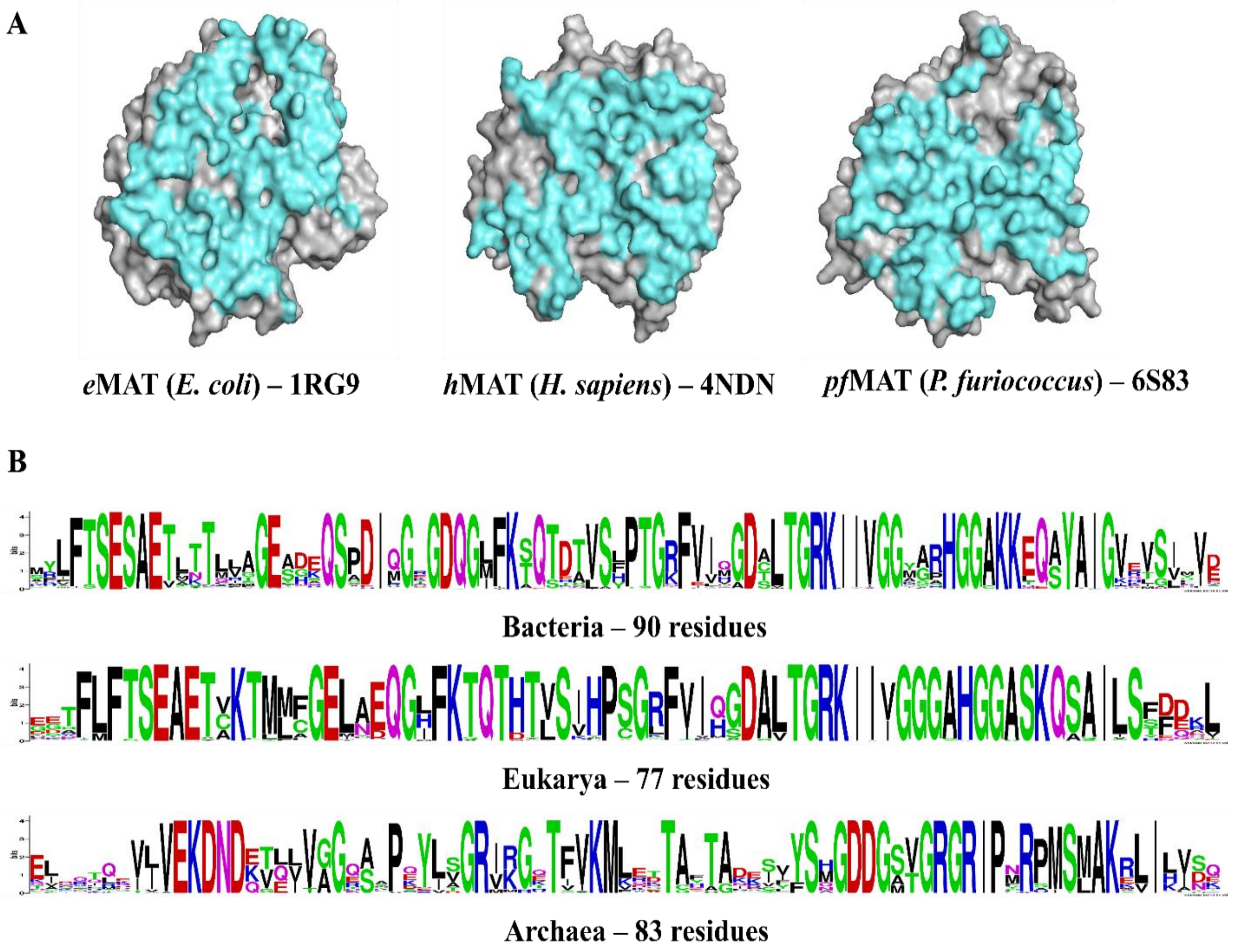


**S1 Fig. (A)** An interface view of three representative MAT structures from the three kingdoms of life with *e*MAT from *E. coli* (PDB: 1RG9), *h*MAT2A from *H. sapiens* (PDB: 4NDN) and *pf*MAT from *P. furiosus* (PDB: 6S83), the interface residues are highlighted in cyan. PISA server was utilized to identify and extract the interface residues [https://www.ebi.ac.uk/pdbe/pisa/].

**(B)** The large interface residues are highlighted as weblogo across the three kingdoms based on sequence alignments performed for 90 interface residues in case of bacteria with corresponding residues for *e*MAT, 77 interface residues for *h*MAT2A and 83 interface residues for *pf*MAT. It is worth mentioning here that the aforementioned residues are only extracted from one chain i.e. chain A and depending on the confirmation and the presence or absence of a ligand – the residue list may change.

### Chains A vs B
### (51 Positions)

**A**

```
E. Coli   (1RG9)                  HLFTEAETYKMLGGEAGQGLFKTDAVFPTDCTGRKGGARGGAKQSAIGTSM
B. pseudomallei (3IML)            YLFTEAETLNLVAGEADQGLFKTDTVLPTDCTGRKGGAPGGAKQSAIGTSM
M. tuberculosis (3TDE)            RLFTEAETLTQHVGEGGQGLFKTDTVLPTDATGRKGGARGGAKQAAIGVGF
C. jejuni (4LE5)                  YLFTEAEVFAKVGGEFNQGIFKTHTVLPTDSTGRKGGSPGGAKQSAIGTSS
T. thermophilus (5H9U)            RLVTEAETLTLFAGEADQGLFKTKTVLPSDTTGRKGGVPGGAKEAAIGVSR
N. Gonorohhea (5T8S)              YLFTEAETLNLVAGEYDQGLFKTDTVLPTDCTGRKGGAPGGAKQSAIGTSS
U. Urealitium (6RKA)              KIITEAEVLCLVAGENNQGIFKSETLISSDTTGRKGGGHGGAKQAAIGVAY
H. Sapiens  (MAT2A - 4NDN)        FLFTEAETVKMLAGEAEQGLFKTHTVHPSDATGRKGGGAGGAKQSAIGLSS
H. Sapiens  (MAT1A - 6SW5)        FMFTEAETVKMLCGEAEQGLFKTHTVHPSDATGRKGGGAGGAKQSAIGLSS
R. norvegicus (1QM4)              FMFTEAETVKMLCGEAEQGLFKTHTVHPSDATGRKGGGAGGAKQSAIGLSS
C. parvum (6C07)                  FLFSEAETCKFMFGEKEQGMFKTHTLLPSDATGRKGGGAGGAKQSGIGLSY
E. hystolytica(3SO4)              FFFTEAETAKLLLGECEQGIFKTHTVYPSDATGRKGGGAGGAKQSAIGLSN
M. truncatula (6VCW)              FLFTEAETCKMMFGENEQGHFKTHTLHPSDATGRKGGGAGGAKQSAIGLSF
A. thaliana (6VCZ)                FLFTEAETCKMMFGENEQGHFKTHTLHPSDATGRKGGGAGGAKQSAIGLSF
P. Furiosus  (6S83)               QVELENQVEVYLSGRRGTSFVKLIDTYTADDVGRGITRHSMAKRLQIGLVS
T. kodakarensis (4L4Q)            KVELENQVEVYLSGRRGTSFVKLIDTYTADDVGRGITRHSMAKRLQIGLVS
S. solfataricus (4K0B)            QVELENKTLVYIAGRKGTSFVKLVDTYTGDDTGRGITRPSLAKQLQIGLIN
                                  #     # ## ##    # #### ## #  ####    # #  ## # #
```

**B**

```
Bacteria_ancestor (model)         YLFTEAETLTLIAGEAGQGMFKTDTVHPTDCTGRKGGGAGGAKQAAIGLSM
E. coli* (1RG9)                   HLFTEAETYKMLGGEAGQGLFKTDAVFPTDCTGRKGGARGGAKQSAIGTSM

Eukarya_ancestor (model)          FLFTEAETAKMMFGEAEQGIFKTDTVHPSDCTGRKGGGAGGAKQSAIGLSF
H. sapiens* (MAT1A - 6SW5)        FMFTEAETVKMLCGEAEQGLFKTHTVHPSDATGRKGGGAGGAKQSAIGLSS

Archaea_ancestor (model)          QVELENQVQVYLSGRRGTSFVKLITTYTADDTGRGITRPSMAKRLQIGLIN
P. furiosus* (6S83)               QVELENQVEVYLSGRRGTSFVKLIDTYTADDVGRGITRHSMAKRLQIGLVS
```

**S2 Fig. (A**) A structural alignment depicting the 51 interface residues from 17 MAT crystal structures obtained from PDB database. Here the organisms name is shown with their respective MAT PDB ID's. In the sequence alignment the conserved positions between bacteria and eukarya MAT are highlighted in red with shared conserved positions from archaea MAT highlighted in blue. Within the alignment the residues which were *not* identified as interface residues by PISA server are highlighted in bold and underlined. We chose a cutoff for including a maximum of five non-interface residues and this data set was utilized to test for the hierarchical clustering (using pvclust package from R language) with charge and hydrophobicity (physiochemical properties). Additionally, we also highlight 24 positions in the alignment that care completely constituted by interface residues and these positions are highlighted by '#' at the bottom of the alignment. A second dataset consisting of these 24 'interface residues only' positions was also utilized to conduct the clustering test (Fig. SI5).

**(B)** A comparison of the 51 positions between representatives of the modern-day MAT from the three kingdoms of life and respective ancestral sequences (from their kingdoms). Here, Archaea ancestral MAT is ArchaeaAnc and the alignment depicts 6 changes in contrast to pfMAT (PDB: 6S83) with eukarya ancestral MAT depicting 8 changes when compared with hMAT1A (PDB: 6SW5) and bacteria ancestral MAT depicting 13 when compared with eMAT (PDB: 1RG9).

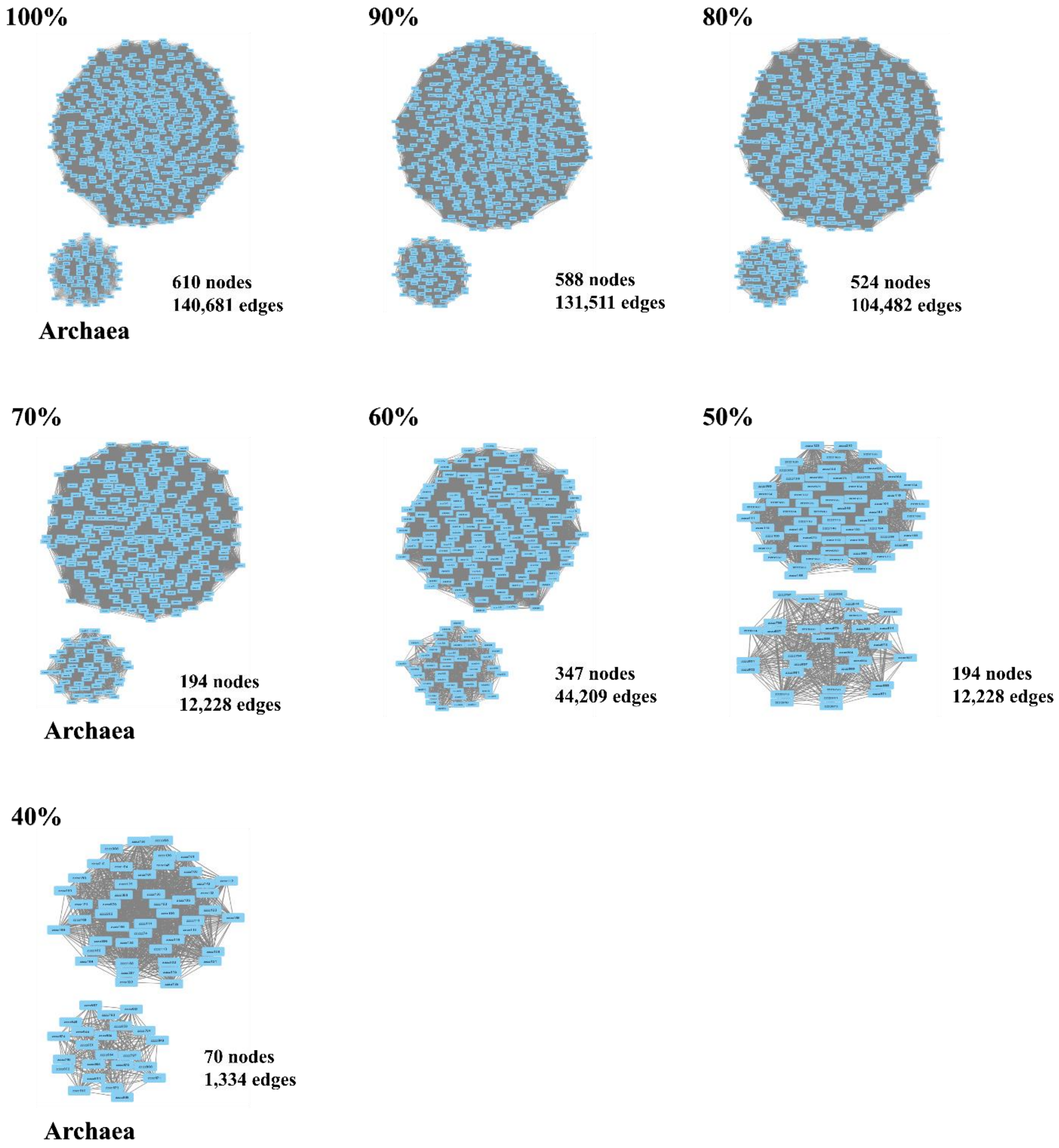


**S3 Fig.** The sequence networks created with EFI tool (post CD-HIT dataset reduction) and visualized at different sequence identity percentage (with default recommendations from EFI web server), visualized with cytoscape. As noted in the figure the two sub-networks form distinct topologies. Each topology (100%-40%) is further annotated with the respective number of nodes and edges.

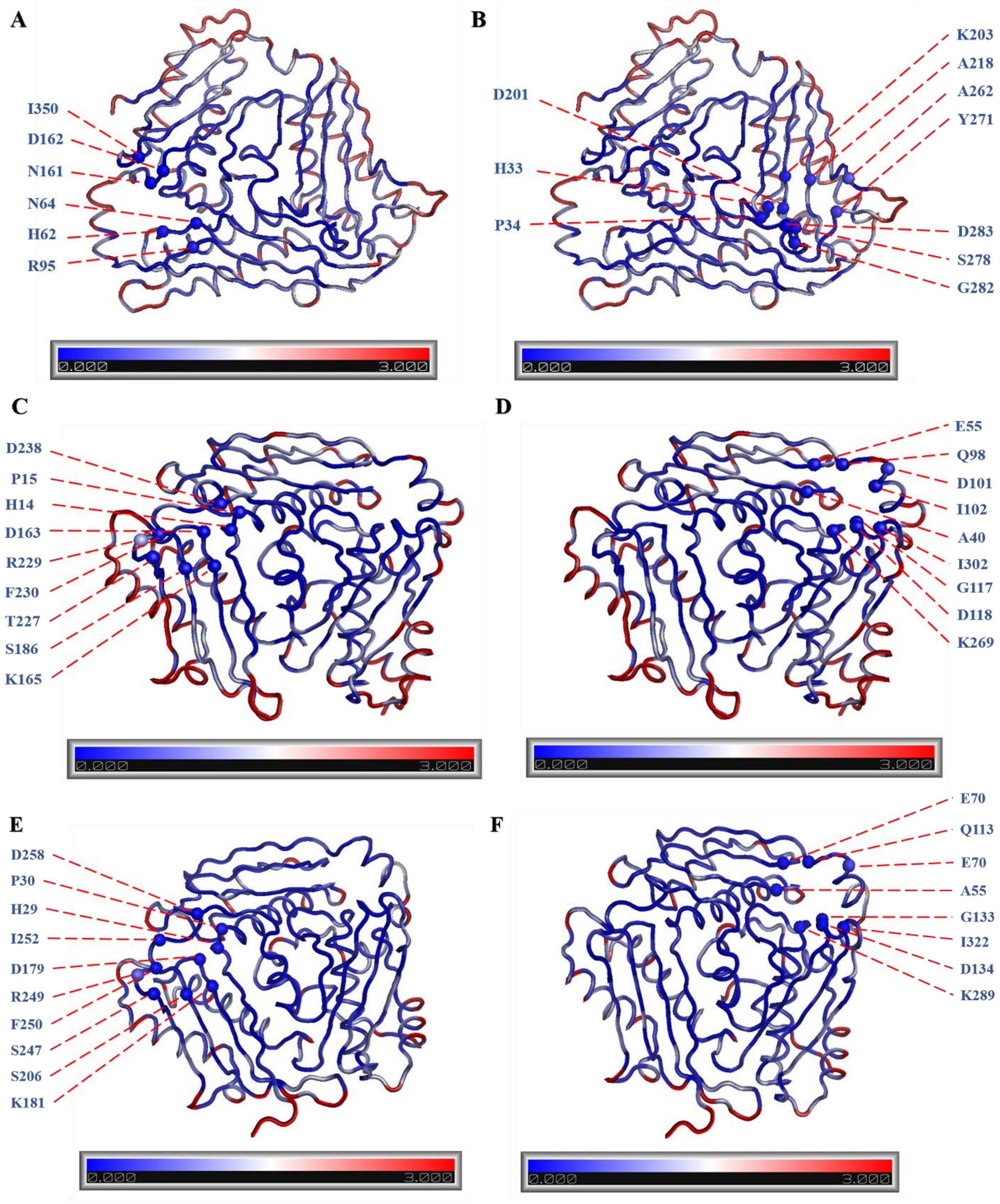


**S4 Fig.** The evolutionary trends adapted by the interface sites (located along the large interface of the homo-tetramer) as well the catalytic sites. Upon normalizing the ML evolutionary rates, we mapped them on to the representative structures (as a function of

B-factors). Here, the rates per-site are designated with a scale wherein the relatively 'slowly' evolving regions are highlighted in blue (~ 0 ML rates), while the sites experiencing 'faster' evolutionary rates are highlighted in red (~3 ML rates). Residues located within a ~4Å of SAM are highlighted in blue and labelled accordingly.

Highlighted here, are the evolutionary rates with an emphasis on the interface region of the tkMAT from archaea *Thermococcus kodakarensis* (PDB: 4l4Q) represented in panels **A** and **B**. The highlighted residues are mentioned below:

*Chain A:* H62, N64, R95, N161, D162, I350
*Chain B:* H33, P34, D201, K203, A218, A262, Y271, S278, G282, D283

eMAT from bacteria *Escherichia coli* (PDB: 1RG9) represented in panels **C** and **D**. The highlighted residues are mentioned below:

*Chain A:* H14, P15, D163, K165, S186, T227, R229, F230, D238,
*Chain B:* A40, E55, Q98, D101, I102, G117, D118, K269, I302

region of the hMAT2A from eukarya *Homo sapiens* (PDB: 4l4q) represented in panels **E** and **F**. The highlighted residues are mentioned below:

*Chain A:* H29, P30, D179, K181, S206, S247, R249, F250, I252, D258,
*Chain B:* A55, E70, Q113, D116, I117, G133, D134, K289, I322

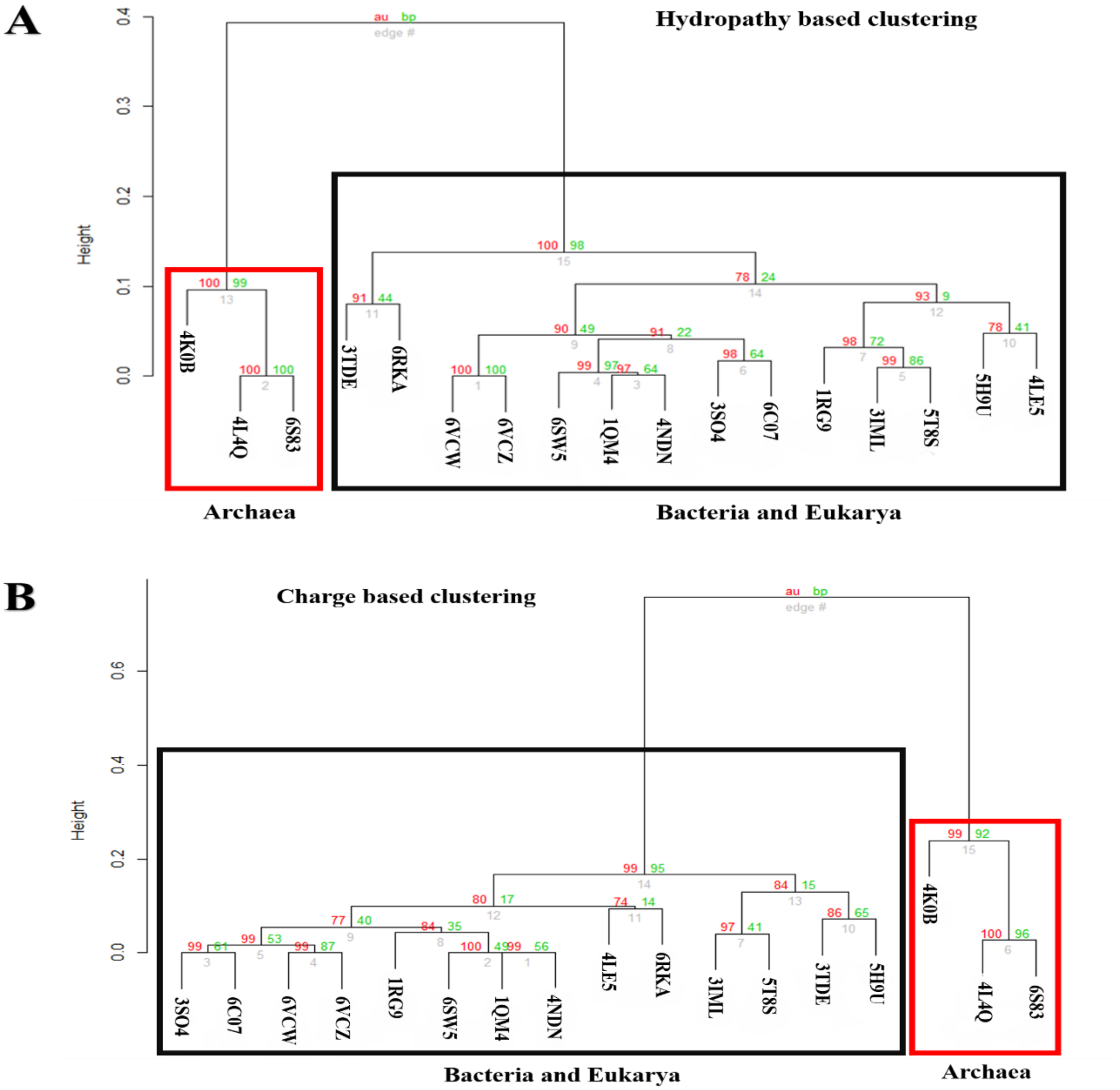


**S5 Fig. Clustering analysis of hydropathy (A) and charge (B) of representative structures from the three kingdoms of life.**

Clustering analysis of the physiochemical properties (Hydropathy and Charge) of 24 structurally aligned large-interface residues from chain A of 17 MAT structures (PDBs list in the SI). Based on the 51 aligned positions we calculated the per-site hydropathy with the help of Kyte-Doolittle scale from the protscale server and per-site charges were calculated with EMBOSS charge server for the aforementioned 17 MAT structures. Subsequently, we conducted clustering analysis with the help of pvclust package in R. This further provides a statistical score in terms of AU (approximately unbiased - in red) p-value and BP (bootstrap probability - in green) value for comparison of the clusters which reveals that the hydropathy **(A)** and charge distribution **(B)** cluster together for bacteria and eukarya MAT structures with archaea MAT structures clustering separately. In both the cases the archaea cluster is provided with high support values with both the AU and BP parameters, 100 % support for a distinct archaea cluster with respect to the two physiochemical properties each.

| | Eukarya d2hj2A1 Domain-I RMSD | %ID | Eukarya d2hj2A2 Domain-II RMSD | %ID | Eukarya d2hj2A3 Domain-III RMSD | %ID |
|---|---|---|---|---|---|---|
| Eukarya d2hj2A1 Domain-I (16-125) | - | - | - | - | - | - |
| Eukarya d2hj2A2 Domain-II (126-251) | 1.86 (L=64) | 22.73 | - | - | - | - |
| Eukarya d2hj2A3 Domain-III (252-395) | 2.21 (L=60) | 22.55 | 2.46 (L=72) | 25.86 | - | - |

| | Bacteria d1rg9A1 Domain-I RMSD | %ID | Bacteria d1rg9A2 Domain-II RMSD | %ID | Bacteria d1rg9A3 Domain-III RMSD | %ID |
|---|---|---|---|---|---|---|
| Bacteria d1rg9A1 Domain-I (1-102) | - | - | - | - | - | - |
| Bacteria d1rg9A2 Domain-II (103-231) | 1.73 (L=62) | 21.82 | - | - | - | - |
| Bacteria d1rg9A3 Domain-III (252-395) | 2.17 (L=62) | 25.68 | 2.50 (L=75) | 18.92 | - | - |

| | Archaea N-terminal Domain-I RMSD | %ID | Archaea Central Domain-II RMSD | %ID | Archaea C-terminal Domain-III RMSD | %ID |
|---|---|---|---|---|---|---|
| Archaea N-terminal Domain-I (Red) | - | - | - | - | - | - |
| Archaea Central Domain-II (Blue) | 1.57 (L=12) | 19.79 | - | - | - | - |
| Archaea C-terminal Domain-III (Cyan) | 1.24 (L=8) | 17.11 | 1.56 (L=15) | 21.88 | - | - |

**S6 Fig. This analysis depicts a comparison of MAT constituent domains within themselves from the same kingdom.**

For instance, within eukarya (PDB: 2HJ2) the three domains d2hj2A1 (16-125), d2hj2A2 (126-251), and d2hj2A3 (252-395) as described in SCOPe database can be structurally aligned with a common length core of 43 residues and an average pairwise RMSD of 3.45Å with A1 and A2 being closer to each other with an RMSD of 1.8Å while A2 and A3 are the closest in terms of sequence identity ~26%. Similar trend is observed in case of bacteria (PDB: 1RG9) with the three domains d1rg9a1 (1-102), d1rg9a2 (103-23) and d1rg9a3 (232-383) and archaea. Here, it is worth mentioning that we were not able to obtain MAT domain demarcation in SCOPe and so we implemented the provided domain demarcation from the literature (PDB: 4L4Q – Schlesier et al., 2013)

| | Bacteria d1rg9a1 Domain-I (1-102) | | Bacteria d1rg9a2 Domain-II (103-231) | | Bacteria d1rg9a3 Domain-III (232-383) | |
|---|---|---|---|---|---|---|
| | RMSD | %ID | RMSD | %ID | RMSD | %ID |
| Eukarya d2hj2A1 Domain-I (16-125) | 0.45 (L = 102) | 62.75 | 1.84 (L = 63) | 15.05 | 2.21 (L = 63) | 25.00 |
| Eukarya d2hj2A2 Domain-II (126-251) | 1.87 (L = 64) | 19.48 | 0.90 (L = 122) | 48.36 | 2.43 (L = 73) | 19.70 |
| Eukarya d2hj2A3 Domain-III (252-395) | 2.23 (L = 61) | 19.48 | 2.43 (L = 69) | 22.95 | 0.87 (L = 143) | 61.11 |

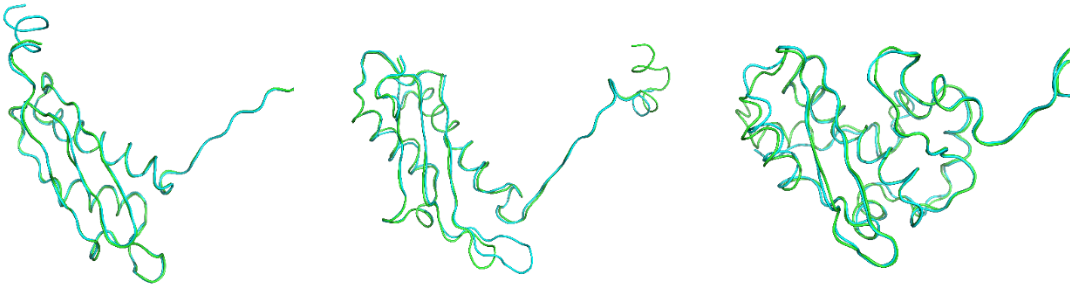

**Domain-I** **Domain-II** **Domain-III**

**S7 Fig. This analysis depicts a comparison of MAT constituent domains across the kingdoms of life.**

For instance, here we compared the three domains from eukarya (PDB: 2HJ2): d2hj2A1 (16-125), d2hj2A2 (126-251), and d2hj2A3 (252-395) as described in SCOPe against their counterparts from bacteria (PDB: 1RG9): with the three domains d1rg9a1 (1-102), d1rg9a2 (103-23) and d1rg9a3 (232-383) and the structural alignment is depicted in the lower panel. This comparison suggests that the three constituent domains clearly share a high level of homology owing to the high sequence similarity and well aligned structures with lower RMSD values. Here, we attempted to align the archaea MAT domains against bacteria and eukarya MAT domains but the low-quality sequence and structural alignment further emphasize the fact that archaea express a highly diverged form of MAT.

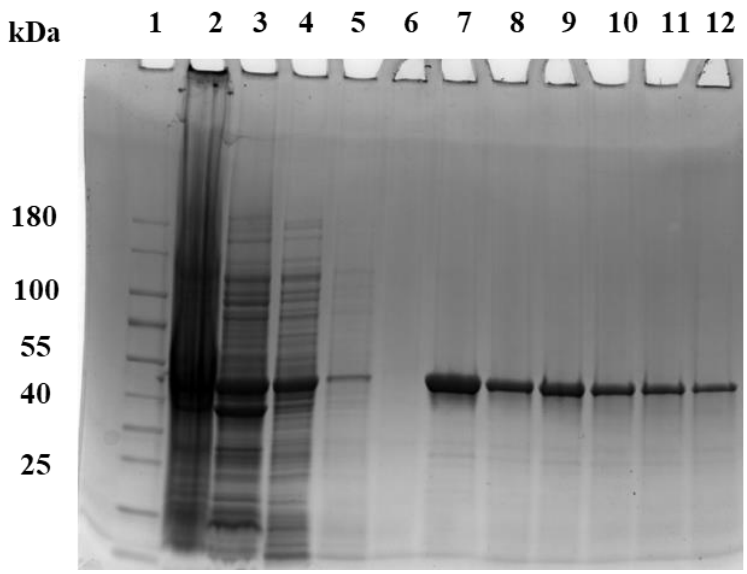


**S8 Fig. SDS–polyacrylamide gel electrophoresis (SDA-PAGE) of ArchaeaAnc MAT protein purification.** 12.5 % SDS–PAGE, Coomassie blue-stained. Lane 1-prestained molecular marker, lane 2- bacterial pellet, lane 3-supernatant, lane 4- flow through, lane 5-wash 1, lane 6- wash 2, lane 7- elution 1, lane 8- elution 2, lane 9- elution 3, lane 10-elution 4, lane 11- elution 5, lane 12-elution 6. Expression and purification as MjMAT.

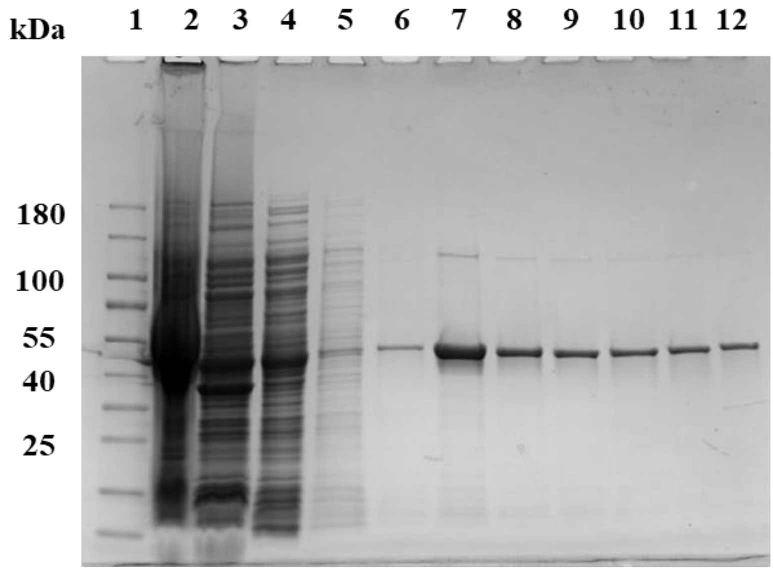


**S9 Fig. SDS–polyacrylamide gel electrophoresis (SDA-PAGE) of CrenAnc MAT protein purification.** 12.5 % SDS–PAGE, Coomassie blue-stained. Lane 1-prestained molecular marker, lane 2- bacterial pellet, lane 3-supernatant, lane 4- flow through, lane

5-wash 1, lane 6- wash 2, lane 7- elution 1, lane 8- elution 2, lane 9- elution 3, lane 10-elution 4, lane 11- elution 5, lane 12-elution 6. Expression and purification as MjMAT.

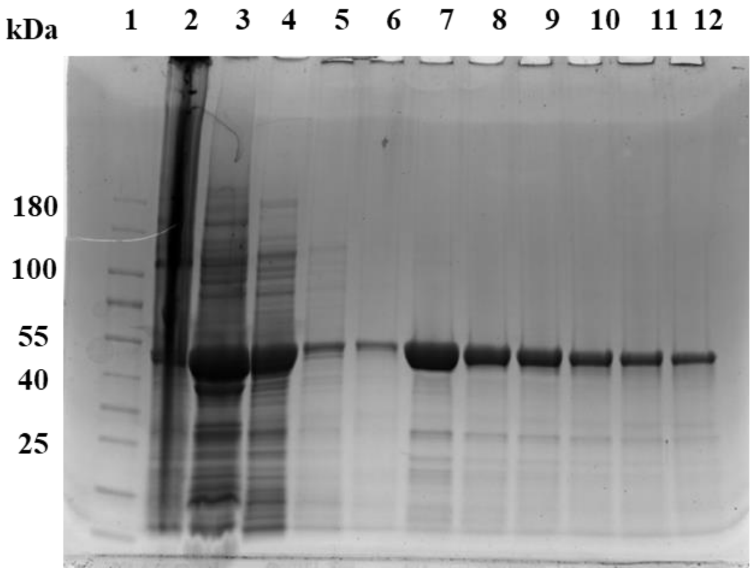


**S10 Fig. SDS–polyacrylamide gel electrophoresis (SDA-PAGE) of EuryAnc MAT protein purification.** 12.5 % SDS–PAGE, Coomassie blue-stained. Lane 1-prestained molecular marker, lane 2- bacterial pellet, lane 3-supernatant, lane 4- flow through, lane 5-wash 1, lane 6- wash 2, lane 7- elution 1, lane 8- elution 2, lane 9- elution 3, lane 10-elution 4, lane 11- elution 5, lane 12-elution 6. Expression and purification as MjMAT.

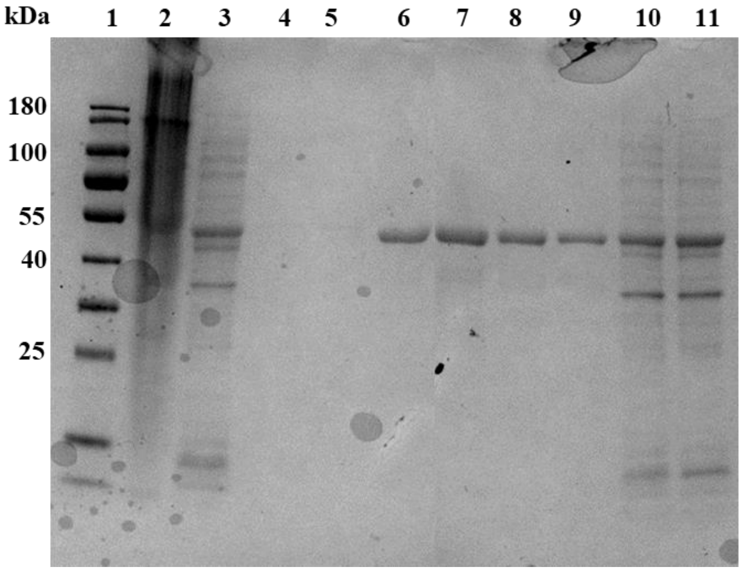


**S11 Fig. SDS–polyacrylamide gel electrophoresis (SDA-PAGE) of MjMAT protein purification.** 12.5 % SDS–PAGE, Coomassie blue-stained. Lane 1-prestained molecular marker, lane 2- bacterial pellet, lane 3- flow through, lane 4-wash 1, lane 5- wash 2, lane 6- elution 1, lane 7- elution 2, lane 8-elution 3, lane 9- elution 4, lane 9- elution 5, lane 10-supernatant, lane 11- cells before sonication. Expression and purification as reported previously[1].

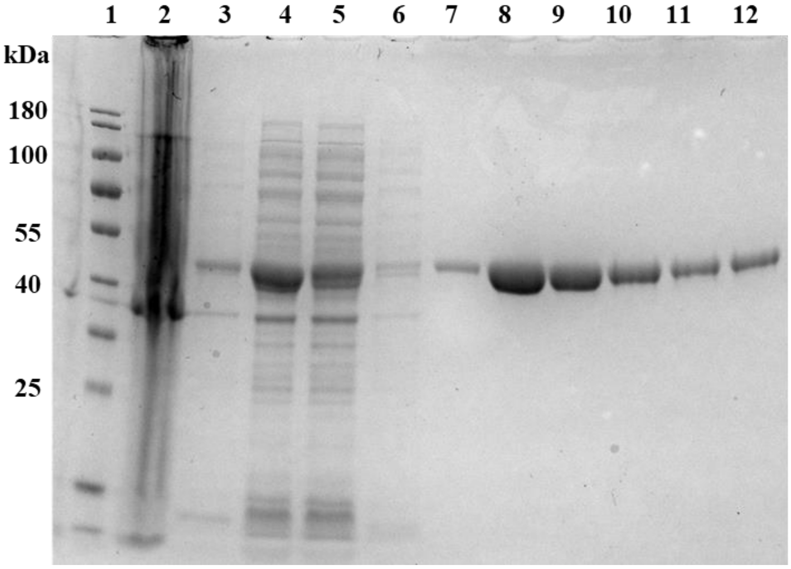


**S12 Fig. SDS–PAGE of hMAT1A purification.** 12.5% SDS–PAGE, Coomassie blue-stained. Lane 1-prestained molecular marker, lane 2- bacterial pellet, lane 3- flow through, lane 4- supernatant, lane 5-wash 1, lane 6- wash 2, lane 7- elution 1, lane 8- elution 2, lane 9- elution 3, lane 10- elution 4, lane 11- elution 5, lane 12-elution 6. Expression and purification as reported previously[2].

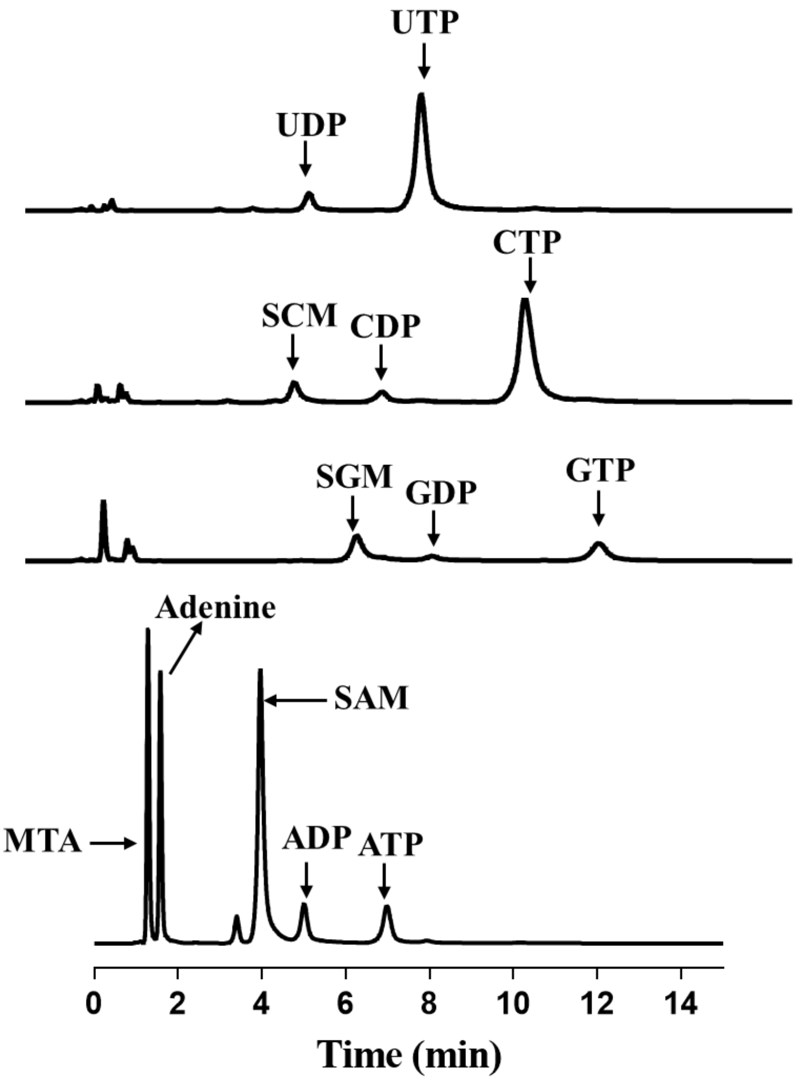


**S13 Fig. UPLC chromatogram of the reaction between NTP, methionine, and ArchaeaAnc.** Reaction details and UPLC method as reported in methods.

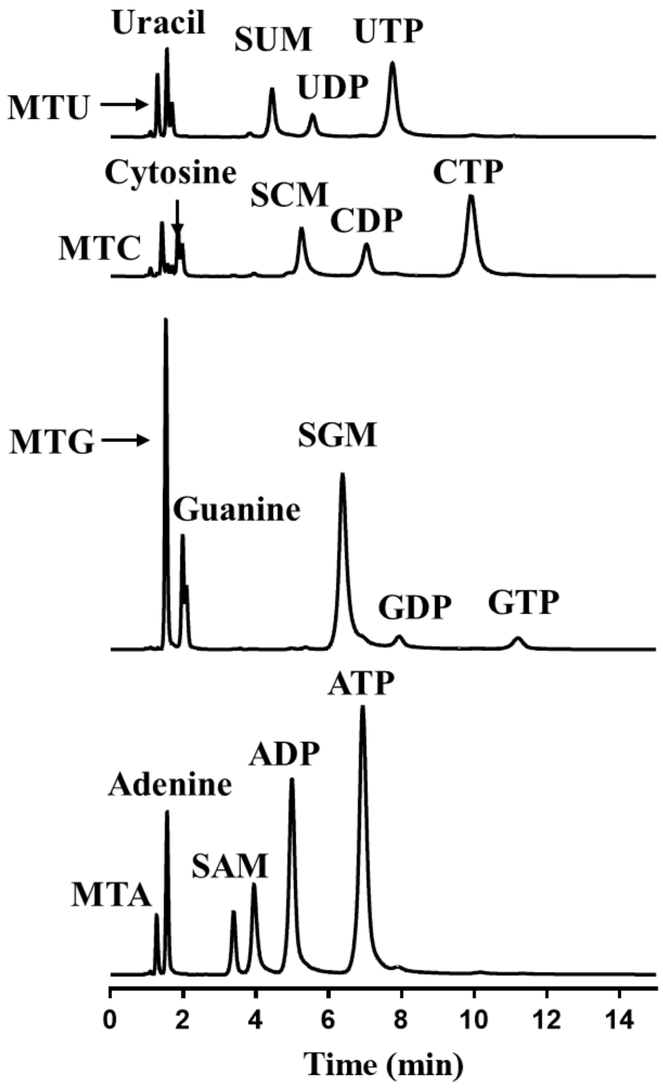


**S14 Fig. UPLC chromatogram of the reaction between NTP, methionine, and CrenAnc.** Reaction details and UPLC method as reported in methods.

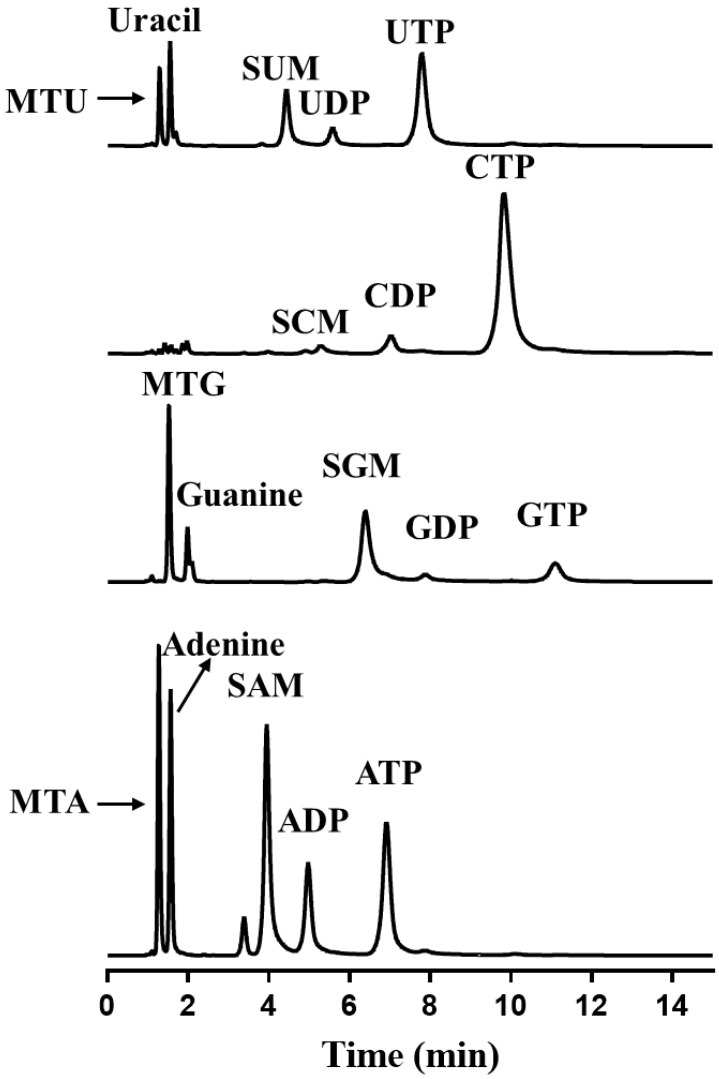


**S15 Fig. UPLC chromatogram of the reaction between NTP, methionine, and EuryAnc.** Reaction details and UPLC method as reported in methods

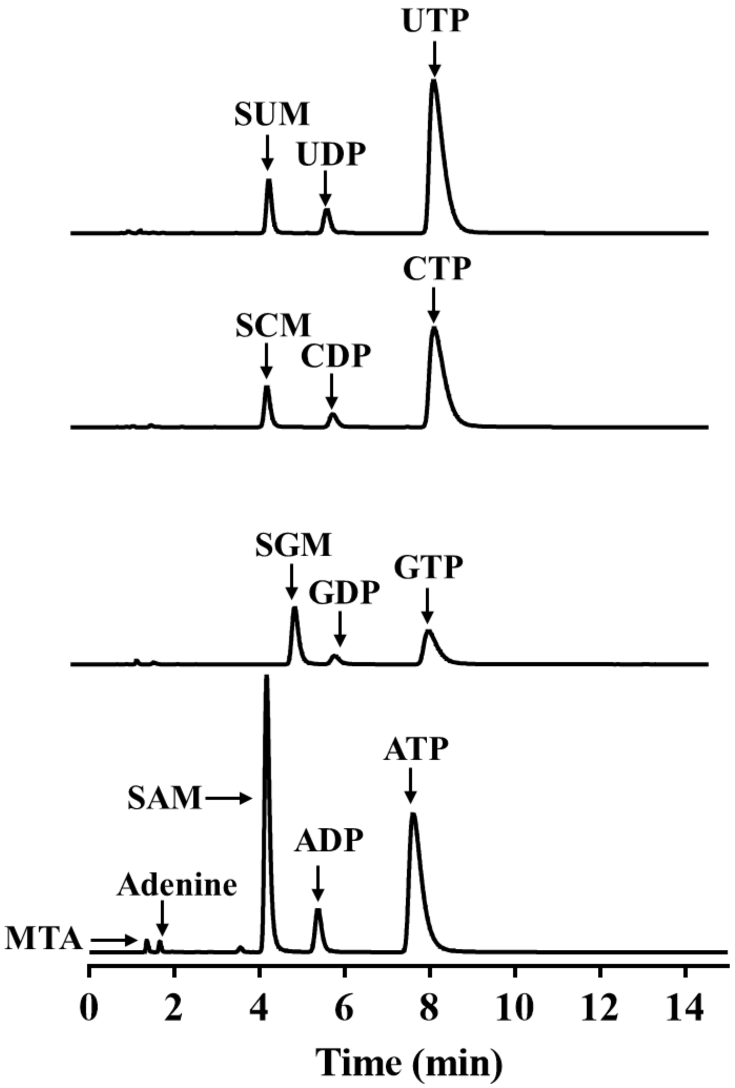


**S16 Fig. UPLC chromatogram of the reaction between NTP, methionine, and MjMAT.** Reaction details and UPLC method as reported in methods.

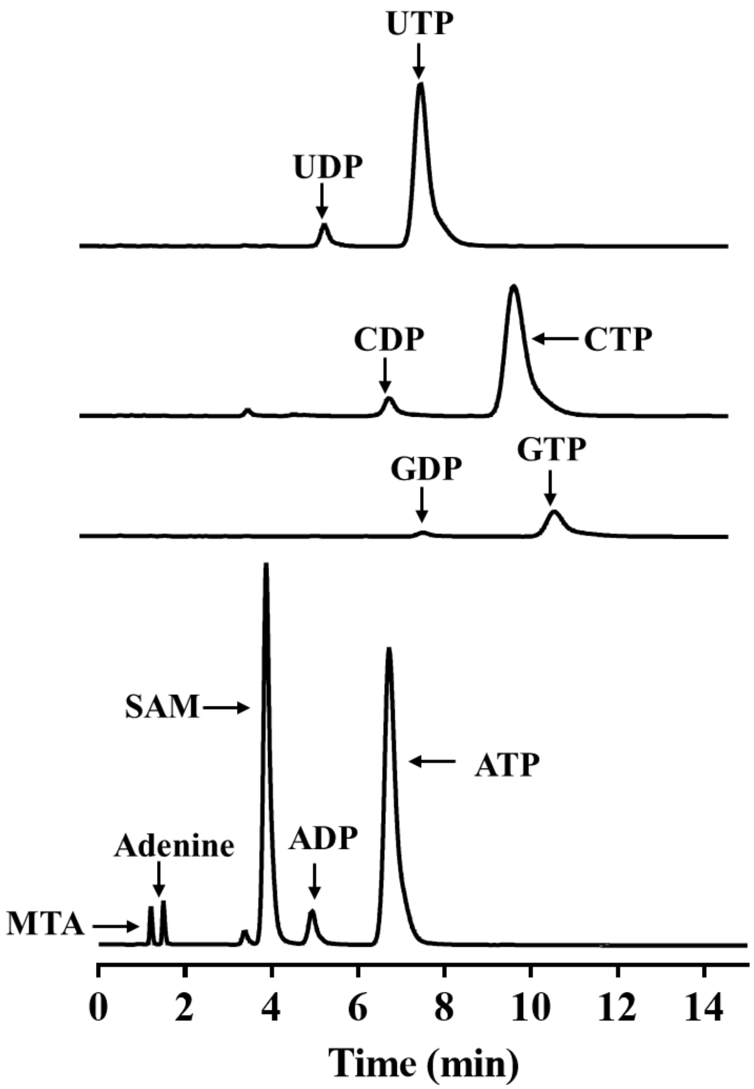


**S17 Fig. UPLC chromatogram of the reaction between NTP, methionine, and hMAT1A.** Reaction details and UPLC method as reported in methods.

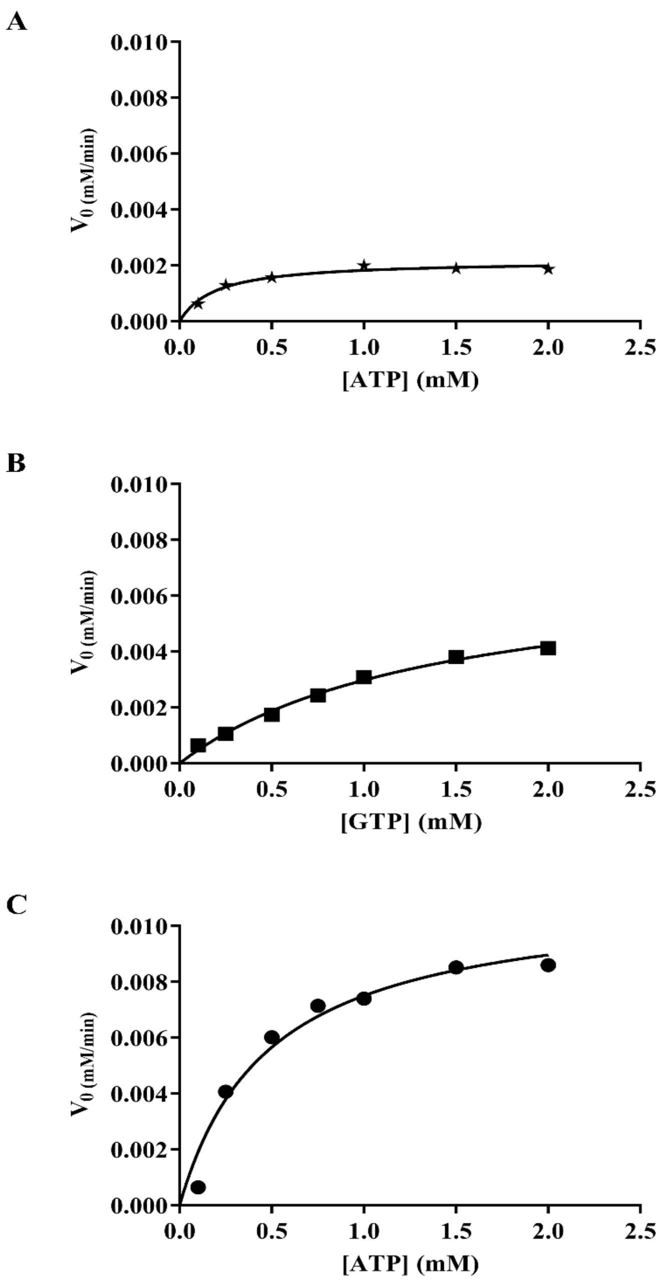


**S18 Fig. Kinetic parameters for the SAM and SGM formation by ArchaeaAnc and hMAT1A.**

Kinetic parameters for the SAM and SGM analog formation by ArchaeaAnc with 0.5 µM using a concentration of ATP (A), GTP (B), in the range of 0.1 to 2 mM and a fixed concentration of methionine (10 mM) in presence of HEPES (100 mM), KCl (50 mM), $MgCl_2$ (10 mM), pH 8 at 55 ℃. Same parameters as of ArchaeaAnc was used for hMAT1A ATP except temperature 37 °C (C). SAM and SGM production was analyzed by UPLC and data fitted to the Michaelis-Menten equation using GraphPad Prism. Experiments were performed in duplicates.
